# *Cis-*regulatory variation in relation to sex and sexual dimorphism in *Drosophila melanogaster*

**DOI:** 10.1101/2022.09.20.508724

**Authors:** Prashastha Mishra, Tania S. Barrera, Karl Grieshop, Aneil F. Agrawal

## Abstract

Much of sexual dimorphism is likely due to sex-biased gene expression, which results from differential regulation of a genome that is largely shared between males and females. Here we use allele-specific expression to explore *cis*-regulatory variation in *Drosophila melanogaster* in relation to sex. We develop a Bayesian framework to infer the transcriptome-wide joint distribution of *cis*-regulatory effects across the sexes. We use this approach to quantify transcriptome-wide sex differences in *cis*-regulatory effects as well as examine patterns of *cis*-regulatory variation with respect to two other levels of variation in sexual dimorphism: (i) across genes varying in their degree of sex-biased expression, and (ii) among tissues that vary in their degree of dimorphism (e.g., relatively low dimorphism in heads vs high dimorphism in gonads). We uncover evidence of widespread *cis*-regulatory variation in all tissues examined, with female-biased genes being especially enriched for this variation. A sizeable proportion of *cis*-regulatory variation is inferred to have sex-specific effects, with sex-dependent *cis* effects being much more frequent in gonads than in heads. Finally, we detect some genes with reversed allelic imbalance between the sexes. Such variants could provide a mechanism for sex-specific dominance reversals, a phenomenon important for sexually antagonistic balancing selection.

## Introduction

Sexual dimorphism is a universal feature of sexually reproducing species, with males and females differing in their appearance, physiology, life history and behaviour (Fairbairn et al. 2007). Males and females carry similar genetic material (other than differentiated sex chromosomes in some species), and these extensive differences in phenotype are largely the result of differential expression of a shared genome (Parsch and Ellegren 2013).

Transcriptional gene regulation is central to the evolution of sex-biased gene expression. Regulation of gene expression can occur through *cis*-acting regulatory elements linked to their target alleles or through *trans*-acting factors. *Cis*-regulatory variants have been reported to be enriched between species relative to their frequency within species in *Drosophila* (Wittkopp et al. 2004; Wittkopp et al. 2008), yeast (Metzger et al. 2017) and mice (Goncalves et al. 2012), among others. This suggests that *cis*-regulatory factors are preferentially used to respond to natural selection for divergent expression. Additionally, *cis*-acting variants are more likely to have additive effects on expression than *trans*-acting variants (Wray 2007; Lemos et al. 2008), which may influence their ability to respond to selection. In contrast, *trans* factors affect expression in multiple genes, and mutations in *trans* may be constrained by pleiotropic side effects (Stern 2000). Lastly, at a practical level, *cis* variants are easier to study. Here, we focus entirely on regulatory variation in *cis*.

Studies indicate that *cis*-regulatory variation is highly pervasive in a wide range of taxa, including humans (Pastinen and Hudson 2004; Stranger et al. 2012), yeast (Kita et al. 2017), mice (Campbell et al. 2008) and *Arabidopsis* (Cubillos et al. 2014). As in other species, *cis-*regulatory variation is common within *Drosophila melanogaster*, with a higher abundance of variation being detected in *cis* than in *trans* (Genissel et al. 2007; Gruber and Long 2009; Osada et al. 2017). Here we examine *cis-*regulatory variation in *D. melanogaster* by measuring differential expression between alleles, termed “allelic imbalance” (henceforth, AI). We investigate its relationship to sexual dimorphism in three ways.

First, we examine whether *cis*-regulatory variation is more common at genes with dimorphic expression (i.e., sex-biased genes). *Cis*-regulatory variation is expected to be enriched in loci undergoing sexual conflict over gene expression. Under the some conditions, sexual conflict can generate balancing selection that stably maintains polymorphisms (Kidwell et al. 1977; Connallon and Clark 2014). If sex-biased genes are more likely to experience sexual conflict than unbiased genes, then one might predict *cis*-regulatory variants to be more common at sex-biased genes. However, even though sexually divergent selection in the past may be responsible for creating the dimorphic expression observed in the present, it is unclear to what extent sexual conflict may persist in genes with sex-biased expression. Sex-biased genes could represent instances of resolved or ongoing conflicts (Bonduriansky and Chenoweth 2009; Rowe et al. 2018). Similarly, unbiased genes could either lack sexual conflict entirely or simply lack the appropriate variation to evolve sex-biased expression to mitigate conflict (Parsch and Ellegren 2013). Based on population genetic analyses of human and fly data, Cheng & Kirkpatrick (2016) concluded that sexual conflict is the most intense in moderately sex-biased genes; this may lead one to predict *cis*-regulatory variation being most common in genes with an intermediate sex bias. Two previous studies of *cis*-regulatory variation in *D. melanogaster* found different patterns from one another and neither matched this prediction (Osada et al. 2017; Puixeu et al. 2023).

Regulatory variants that affect expression similarly in both sexes would not allow for the resolution of sexual conflict via sex-biased gene expression. Thus, our second goal is to examine the extent to which *cis*-regulatory effects differ between the sexes, measured as ‘sex-dependent’ allelic imbalance. In terms of the transcriptome-wide joint distribution of *cis*-regulatory effects across the sexes, we can consider sex differences from two related perspectives: the frequency with which AI differs between sexes or the intersexual correlation in AI. The latter harkens to the intersexual genetic correlation in expression (often represented as *r_MF_*), which governs the extent to which expression can evolve independently between the sexes.

The most extreme form of sex-dependent AI occurs when there is a reversal of the dominant allele in the sexes (“sex-reversed AI”), whereby heterozygous males have >50% expression from one allele, while heterozygous females have >50% expression from the alternative allele. We perform additional analyses to find instances of sex-reversed AI. Cases of sex-reversed AI hold interest because they provide a potential mechanism for sex-specific dominance reversals in fitness, whereby heterozygotes of each sex could dominantly express the preferred allele for a locus under sexually antagonistic selection. Theoretical models indicate that sexually antagonistic variants exhibiting dominance reversals are preferentially maintained by balancing selection, due to partially mitigating sexual conflict (Kidwell et al. 1977; Spencer and Priest 2016; Connallon and Chenoweth 2019; Grieshop et al. 2024).

Our third goal was to examine *cis*-regulatory variation across tissues varying in their degree of sexual dimorphism. We studied expression data from whole body, head, and gonad samples. Gonads and heads offer an interesting comparison because gonads are highly sexually dimorphic whereas heads are much less so. We compare the frequency of AI and particularly, sex-dependent AI between these two tissues to discern if patterns in AI reflect the underlying levels of sexual dimorphism.

## Methods

### Sample preparation and sequencing

We crossed two inbred lines of *D. melanogaster* to obtain F_1_ individuals for the study. DGRP-177 belongs to the Drosophila Genetic Reference Panel, a suite of genotypes derived from natural populations of *D. melanogaster* in North Carolina (Mackay et al. 2012). SP159N was derived from a South African population, as part of the Drosophila Population Genomics Project (Pool et al. 2012; Lack et al. 2016). We crossed DGRP-177 males with SP159N females (“main cross”), and also performed a reciprocal cross. All flies were reared and maintained at 25°C on a 12-hr light-dark cycle. After eclosion, virgin F_1_ males and females were collected and kept in separate vials for 3 days. Vials that were observed to have eggs laid in them were discarded. The F_1_ individuals from the main cross (DGRP-177 males × SP159N females) were used to obtain samples of whole body flies, heads, and gonads, with three replicate samples of each sex*tissue-type combination. From the reciprocal cross, only whole body samples were collected (three replicates of each sex). Testes and ovaries were obtained by dissection of anaesthetised whole flies, performed over ice, with the flies placed in PBS buffer. Each replicate for gonads consisted of 10-12 individuals, while each replicate for whole flies and heads consisted of 5-7 individuals. Heads and whole flies were flash-frozen in liquid nitrogen and then stored at −80°C as they awaited RNA extraction; gonads were stored at −80°C upon dissection. RNA extraction was performed using ThermoFisher PicoPure RNA Isolation Kit. Paired-end sequencing at a read length of 200 bp was performed using Illumina NovaSeq6000. In addition, DNA was extracted from the parents and F_1_ individuals using the Qiagen DNeasy Blood and Tissue Kit. Paired-end whole genome sequences were obtained at a read length of 100 bp using Illumina HiSeqX.

### Genotype-specific references

Genotype-specific references are constructed from the reference genome by incorporating single nucleotide polymorphisms (SNPs) for each parental genotype separately to reduce mapping bias when analysing allele-specific expression (Rozowsky et al. 2011; Graze et al. 2012). We called variants (SNPs and indels) for both DGRP-177 and SP159N using the GATK Best Practices workflow (McKenna et al. 2010; Auwera et al. 2013). The whole genome sequences for DGRP-177 and SP159N were aligned to the *D. melanogaster* Release 6 reference genome (dos Santos et al. 2015) using BWA (Li and Durbin 2009) with default parameters. The resulting alignment file was processed using GATK tools to sort reads and mark PCR/optical duplicates. Variants were called by using GATK HaplotypeCaller and the resulting VCF file was separated into two VCFs, one each for indels and SNPs. We obtained a BED file of coordinates corresponding to heterozygous SNP calls and regions around indels for each parental genome. Depending on the length of the indels, the following coordinates were included in the BED file: 10 bp around indels of length ≤ 6 bp, 20 bp around indels of length > 6 bp but ≤ 12 bp, and 100 bp around indels > 12 bp. The coordinates were then masked in the reference genome with N’s. We masked sites without high certainty of a homozygous SNP call in both parents, i.e., any sites where the major allele was < 95%. We also masked sites where the depth of coverage fell below 10. Following recommendations by GATK, the remaining variants—the homozygous SNPs—were subject to hard-filtering based upon quality measures (Table S1). The filtered SNPs corresponding to DGRP-177 and SP159N were incorporated separately into the masked reference genome, to create genotype-specific references for each parent.

### Competitive mapping

For each sample, RNA-seq reads were aligned to both genotype-specific references using STAR v2.7 (Dobin et al. 2013) with default parameters. Following a similar method to Sánchez-Ramírez & Cutter (2021), a Python-based competitive read mapping approach was used to obtain allele-specific read counts from RNA-seq data for the F_1_ heterozygotes. The two BAM alignments yielded by STAR were sorted and indexed by read names. The alignment score (AS) and number of mismatches (nM) of each read when aligned to the two references were used to assign each read’s origin as follows: a) if a read had a higher AS when aligned to a given parent’s genotype-specific reference, it was ascribed as belonging to that genotype, b) if the read has equal AS for both alignments, the read was assigned to the genotype that yielded a lower nM, and c) if AS and nM do not differ between the two alignments for a read, it was assigned to neither genotype, i.e., an “ambiguous” read. (We also tried the competitive alignment using only nM to ascribe parental origin and obtained results similar to that from using both AS and nM.) Ambiguous reads were excluded from further analysis. The percentage of such informative (i.e., non-ambiguous) reads ranged from 18% to 43% for the samples examined (Table S2). Competitive read mapping yielded two alignment files, corresponding to allele-specific reads of each parent. Finally, allele-specific read counts were obtained from those BAM alignments using HTSeq-count v0.12 (Anders et al. 2015), with secondary or chimeric alignments being ignored when aggregating reads.

Bias in mapping can occur for multiple reasons and lead to errors in estimation of allele-specific expression (Degner et al. 2009; León-Novelo et al. 2014). Genomic DNA from F_1_s should have 50% of reads mapping to each parent in the absence of mapping bias. As described in the Supplementary Material, we used genomic DNA from male and female F_1_ samples to conservatively filter out genes with possible mapping bias (Supplementary Text S1, Figures S1-S3, Tables S3-S5).

### Parental effects in AI

Parental effects on allele-specific gene expression in *Drosophila* are believed to be rare (Wittkopp et al. 2006; Coolon et al. 2012; Puixeu et al. 2023). Still, certain forms of mapping bias could manifest as apparent parental effects. To detect real or apparent “parental” effects, and then remove such genes from further analyses, we applied generalised linear models to the whole body samples from the main and reciprocal crosses utilising the R package *lme4* v.1 (Bates et al. 2007). The first test was applied separately to each sex. Only genes that had read counts >30 in all the 6 replicates were considered. For each gene, we analysed the proportion of reads assigned to the DGRP-177 genome with a quasibinomial model, including cross direction (main versus reciprocal) as a fixed effect. The intercept term of this model indicates allelic imbalance in the absence of an effect of cross direction. Additionally, we analysed the data for both sexes in a single model, including sex and “cross direction” as fixed effects, but without an interaction term. This test was applied to all genes with a total read count >20 in all 6 male and female replicates and average read count >30. All genes that had a significant cross-direction effect (p < 0.05) were excluded from subsequent analyses.

### Inference of allelic imbalance and sex differences in allelic imbalance

For our analyses examining AI and sex-depenent AI (hereafter “SD-AI”), the data for each “tissue” type (gonads, heads, whole bodies) were analysed separately. Whole bodies from the main and reciprocal crosses were also analysed separately for two reasons. First, as we expect similar patterns in the whole body analyses from each cross direction, any major discrepancies could serve as a cause for concern. Second, the whole body results are comparable to results for gonads and heads because all the analyses are based on three replicates per sex for each tissue type. Only genes with at least 30 assignable reads in each of the replicates were included in the analysis for a given tissue. For each gene, we analysed the proportion of reads assigned to the DGRP-177 genome with a quasibinomial model, including sex as a fixed effect and using sum contrasts. In this model, the intercept term indicates the sex-averaged AI effect and the “sex” term indicates the difference in AI between sexes (i.e., a significant “sex” term is evidence of sex-dependent AI).

### Inference of tissue effects on allelic imbalance

We performed analyses contrasting heads and gonads, parallel to our analyses of the two sexes. Separately for each sex, we first analysed the gonad and head data with a quasibinomial model including a tissue term. Significance of the intercept term is taken as evidence of (tissue-averaged) AI. A significant tissue term (p < 0.05) indicates tissue-dependent AI. Unexpectedly, we noted substantial differences in the frequency of AI and tissue-dependent AI in the male and female analyses. Subsequently, we examined those genes for which there was sufficient data in both sexes using a quasibinomial model that included sex, tissue, and their interaction. The interaction term indicates sex differences in tissue-dependent AI.

### Inference of sex-and tissue-reversed allelic imbalance

The strongest form of sex-dependent AI—sex-reversed AI—occurs when one allele is responsible for greater than 50% expression in males, while the other allele is more highly expressed in females. To maximize our power, we jointly analysed the whole body data from the main and reciprocal crosses. All genes with read counts <20 in any of the replicates were excluded from analysis. Additionally, we excluded genes where the average read count across all the 12 replicates was <30. We then tested each sex separately for AI, using a quasibinomial model that only fit an intercept term, which signified the AI effect. A gene was said to have sex-reversed AI if it fulfilled the following conditions: (i) it had a significant AI effect in females, (ii) it had a significant AI effect in males, and (iii) the AI effect in the sexes were in opposite directions. The number of false positives expected from this procedure depends on the assumptions one makes for a null hypothesis. We considered several possibilities to arrive at a conservative estimate of the number of false positive cases of dominance reversal (Supplementary Text S1).

Likewise, we used data from gonads and heads to detect cases of tissue-reversed AI in males and females separately. Again, after excluding genes with low read counts as above, we tested each tissue separately for AI using a quasibinomial model where the intercept term represented AI. A gene was said to have tissue-reversed AI if it showed significant AI in both heads and gonadal tissue, and the direction of the AI was opposite between the two tissues. The number of false positive cases of tissue-reversed AI were estimated in a manner analogous to that for sex-reversed AI (Supplementary Text S1).

### Modeling the distribution of AI and SD-AI

In the earlier section, we described analyzing each gene individually to test for the presence of AI and SD-AI. As with any set of frequentist analyses, the results will be subject to both false positives and false negatives. Moreover, estimates of the average magnitudes of AI and SD-AI using “significant” genes will be subject to bias due to the “winner’s curse”. As an alternative, we used a Bayesian approach to model the joint distribution of AI effects across sexes, separately for each tissue.

As the true features of this distribution are unknown, there many ways one could choose to model it. Our goal was to approximate the distribution using relative simple models framed with respect to our key biological interests. To this end, we developed two models, each characterized by four parameters. The first model framework characterizes the distribution in terms of (i) the frequency of genes with non-zero sex-averaged AI, *F_AI_*; (ii) the standard deviation in sex-averaged AI, *σ_AI_*; (iii) the frequency of genes with SD-AI among those with non-zero sex-averaged AI, *F_SD-AI_*; and (iv) the standard deviation in the sex difference in AI, *σ_SD-AI_*. We model AI as a deviation in expression of allele 1 from 50% and sex differences in AI as the difference of allele 1 expression in males from that in females. Because we assume there is no net directionality to AI or SD-AI effects (i.e., there is no bias across genes, on average, of which parental strain allele is more highly expressed), the average values of AI and SD-AI are zero; thus, *σ_AI_* and *σ_SD-AI_* represent the average magnitudes of AI and SD-AI, respectively. The second model framework is similar but instead characterizes the distribution in terms of (i) the frequency of genes with non-zero sex-averaged AI, *F_AI_* (as in the first model framework); (ii) the standard deviation in AI in females, *σ_AI,F_*; (iii) the standard deviation in AI in males, *σ_AI,M_*; and (iv) the intersexual correlation in AI effects, *ρ_MF_*. The first model framework is motivated primarily by an interest in estimating the fraction of AI effects that differ between the sexes (*F_SD-AI_*) whereas the second is motivated primarily by a related but different property, the intersexual correlation in AI effects, *ρ_MF_*.

For both model frameworks, the likelihood function relating the observed read counts to the distribution incorporates a fifth (nuisance) parameter to allow for overdispersion. Because genes for which we have a high amount of data (e.g., highly expressed genes) could have a disproportionate influence on the inferred distribution of AI effects, we performed the analysis on a filtered data set for each tissue type. We removed genes in the bottom quartile of total read count for either sex and then down-sampled the read counts to equal levels for all genes (separately by sex). It is worth remembering that this (and other) filtering steps non-randomly excludes genes from analysis (e.g., genes with low expression in either or both sexes) and, thus, the inferred distribution may not be representative of such genes. We used the R package *BayesianTools* (Hartig et al. 2023) to estimate a posterior distribution. Further details about the models are provided in the Supplementary Text S3.

### Differential gene expression analysis

We obtained gene-level read counts by using *htseq*-*count* on BAM alignments of each RNA-seq sample to the *D. melanogaster* reference genome. We then estimated differential gene expression between the sexes using the R package *DESeq2* (Love et al. 2014). In estimating differential expression, we removed any genes averaging fewer than 50 reads across all the replicates for a given tissue. Differential expression at each gene is estimated as the log_2_ fold change in male-to-female gene expression (log_2_FC). Excluding a small number of genes where the log_2_FC had a standard error greater than 1.5, all other genes were assigned to one of five sex-bias bins, as follows: highly female-biased (-∞ < log_2_FC ≤ −2), moderately female-biased (−2 < log_2_FC ≤ −0.5), unbiased (−0.5 ≤ log_2_FC ≤ 0.5), moderately male-biased (0.5 < log_2_FC ≤ 2), highly male-biased (2 < log_2_FC < ∞). Genes with relatively high uncertainty (i.e., standard error for log_2_FC > 1.5) were not assigned to any of these bins. The four tissue types—heads, gonads, whole bodies (main cross) and whole bodies (reciprocal cross) —were separately analysed for sex-biased expression.

## Results

We crossed two inbred lines of *Drosophila melanogaster* (DGRP-177 and SP159N) and measured allele specific expression in heads, gonads, and whole bodies of F_1_s of both sexes. For whole body samples, we had F_1_s from both cross directions. A series of filters based on parental and F1 genomic data were applied to remove genes with possible mapping biases (Methods).

### Apparent signals of parental effects

Studies suggest that parental effects on allelic expression rarely occur in *Drosophila melanogaster* (Wittkopp et al. 2006; Coolon et al. 2012; Chen et al. 2015). Moreover, patterns consistent with parental effects may be due to residual mapping bias. We performed three tests using whole body samples from the main and reciprocal crosses to identify genes with evidence of “parental effects,” seeking to remove these genes from subsequent analysis. We analyzed the data from each sex separately, and also performed an analysis using combined data from the sexes. 346 out of 7716 (4.5%; X genes included) genes tested in females and 829 out of 8817 (9.4%; X and Y genes excluded) genes tested in males showed a significant effect of cross direction on allelic imbalance (p < 0.05). (The substantially higher incidence of “parental effects” in F_1_ males relative to females could be due to males of the two crosses having *trans*-regulatory differences attributable to complementary XY combinations.) In the combined data from both sexes, 532 of 6371 (8.4%; X and Y genes excluded) genes had a significant effect of cross direction. A total of 10522 genes were analyzed across these three tests, with 1375 (13.1%) showing evidence of parental effects in at least one of the tests. While many of these genes are expected to be false positives (5% of all genes in each test), our primary goal here is to reduce the possibility of mapping bias influencing our analyses of AI. We therefore excluded all these genes showing significant parental effects from further analysis.

### Sex-dependent effects on allelic imbalance

Allelic imbalance appears to be pervasive in the *D. melanogaster* transcriptome, with more than 30% of the genes tested for each tissue showing significant AI (Table 1). Given our *p*-value threshold, we expect ∼5% of genes to falsely appear as showing significant AI (i.e., false positives); the observed fraction is much greater than expected under the null hypothesis. We use a *p*-value criterion for the ease of comparisons among tissues, which would be complicated when applying a false discovery rate; false discovery rate *q*-values depend not only on the data from the focal genes but also the distribution of *p*-values within a group, which could cause genes with similar evidence for AI to have different *q*-values in each group. Nevertheless, a version of Table 1 using FDR cut-offs is shown in Table S8. As expected, using the more conservative FDR requirement reduces the overall frequency of detection of AI but the main patterns are unchanged.

**Table 1.** Frequency of AI and sex-dependent AI.

| Tissue | Number of genes tested | % genes with AI | % genes with sex-dependent AI |
| --- | --- | --- | --- |
| gonads | 2547 | 31.6% | 25.3% |
| heads | 4062 | 45.0% | 10.1% |
| whole bodies (main) | 3627 | 36.5% | 19.6% |
| whole bodies (reciprocal) | 3901 | 35.7% | 18.3% |

In each tissue, numerous genes exhibit significant sex-dependent allelic imbalance, SD-AI (Table 1). About 25% of genes exhibited significant SD-AI in gonads, the most sexually dimorphic tissue in our dataset. In contrast, only 10% of genes show SD-AI in heads. We considered the possibility that our inferences of AI or SD-AI may be a consequence of mapping bias, despite our efforts to minimize such errors. If mapping bias persists in the dataset despite the filters, we would expect patterns in genomic data to be reflected in RNA-seq. Firstly, we would expect a positive correlation in fraction of DGRP-177 reads between F1 genomic and F1 transcriptomic datasets in both the sexes. However, the correlations are very close to zero (Figures S2 & S3; Tables S4 & S5), indicating that mapping bias is unlikely to be a widespread problem among genes inferred to have AI. Secondly, we would expect a significant positive correlation in the sex difference in the fraction of DGRP-177 reads between genomic and transcriptomic datasets. These correlations are also close to zero, remaining so even as we consider only genes with significant sex-dependent AI (Figure S4; Table S6). Thus, our inferences of AI or SD-AI are unlikely to be substantially marred by mapping bias.

### Estimating the distribution of AI and SD-AI

The preceding section summarizes the results from assessing the statistical significance of each gene individually. As such, Table 1 reflects the frequencies of genes for which there was statistical power to detect evidence of AI and SD-AI. Thus, the values in Table 1 not only depend on the aspects of biology that are our real interest (frequency and effect sizes of AI and SD-AI), but also the number of samples, sequencing depth, and the choice of statistical threshold for significance. As an alternative to the approach used to generate Table 1, we employed a Bayesian analysis to model the underlying joint distribution of AI across the sexes in each tissue type. The results are summarized in Table 2. Across all tissues, the inferred frequency of genes with AI was similar (*F_AI_* = 38-48%) as was the magnitude of the (sex-averaged) effect (∼5%). However, there were major differences between tissues with respect to the frequency of sex-differences in AI. In gonads, almost all genes with AI were inferred to have sex differences (*F_SD-AI_* = 94%), whereas in heads very few were (*F_SD-AI_* = 5%). In gonads, the effect sizes of SD-AI tend to be quite large relative to the sex-average effect size of AI. (This is related to a result shown in a later section: AI effects tend to be larger in male than female gonads.) The values in Table 2 represent, to our knowledge, the only available estimates of the distribution of AI/SD-AI effects but, as with any statistical model, some caution is warranted as these estimates are contingent on the assumptions underlaying the framework of the modelled distribution.

**Table 2.** Parameter estimates [with 95% high posterior density interval] for the joint distribution of AI effects across sexes.

| | Frequency of genes with AI, $F_{AI}$ | Average magnitude of sex-averaged AI<sup>1</sup>, $\sigma_{AI}$ | Frequency of genes with SD-AI among genes with AI, $F_{SD-AI}$ | Average magnitude of sex difference in AI<sup>2</sup>, $\sigma_{SD-AI}$ | Overdispersion parameter, $\nu$ |
| --- | --- | --- | --- | --- | --- |
| Gonads | 0.48<br>[0.45, 0.52] | 0.053<br>[0.0496, 0.056] | 0.94<br>[0.90, 1.00] | 0.105<br>[0.099, 0.114] | 0.006<br>[0.005, 0.006] |
| Heads | 0.42<br>[0.40, 0.45] | 0.050<br>[0.054, 0.060] | 0.05<br>[0.01, 0.08] | 0.074<br>[0.051, 0.106] | 0.001<br>[0.001, 0.001] |
| Whole Body (main) | 0.45<br>[0.44, 0.47] | 0.049<br>[0.048, 0.051] | 0.75<br>[0.71, 0.79] | 0.059<br>[0.058, 0.061] | 0.001<br>[0.001, 0.002] |
| Whole Body (reciprocal) | 0.38<br>[0.35, 0.40] | 0.055<br>[0.053, 0.058] | 0.66<br>[0.60, 0.71] | 0.065<br>[0.060, 0.070] | 0.003<br>[0.002, 0.003] |

<sup>1</sup> $\sigma_{AI}$ is the standard deviation of sex-averaged effects, which is modelled as approximately normally distributed with mean 0.
<sup>2</sup> $\sigma_{SD-AI}$ is the standard deviation of sex differences in AI effects, which is modelled as approximately normally distributed with mean 0.

### AI in relation to sex-biased gene expression

We examined how the patterns of AI vary with the degree of sex bias in expression by fitting a Bayesian model separately to gene sets binned by sex bias. For this section, we used an alternative framework for the distribution of AI effects, parameterizing the distribution in terms of the frequency with which AI occurs, *F_AI_* (as in the previous section), as well as the correlation of AI effects between males and females, *ρ_MF_*. There is heterogeneity in the frequency of AI with respect to sex bias (Fig. 1A; Tables S10-12). In heads, *F_AI_* declines from female-biased to unbiased to male-biased genes. There is a similar trend in the whole body samples, though here it is noteworthy that the inferred average effect sizes also vary with lower estimates for female-than male-biased genes (Fig. 1A; Table S10-11).

**Figure 1.**
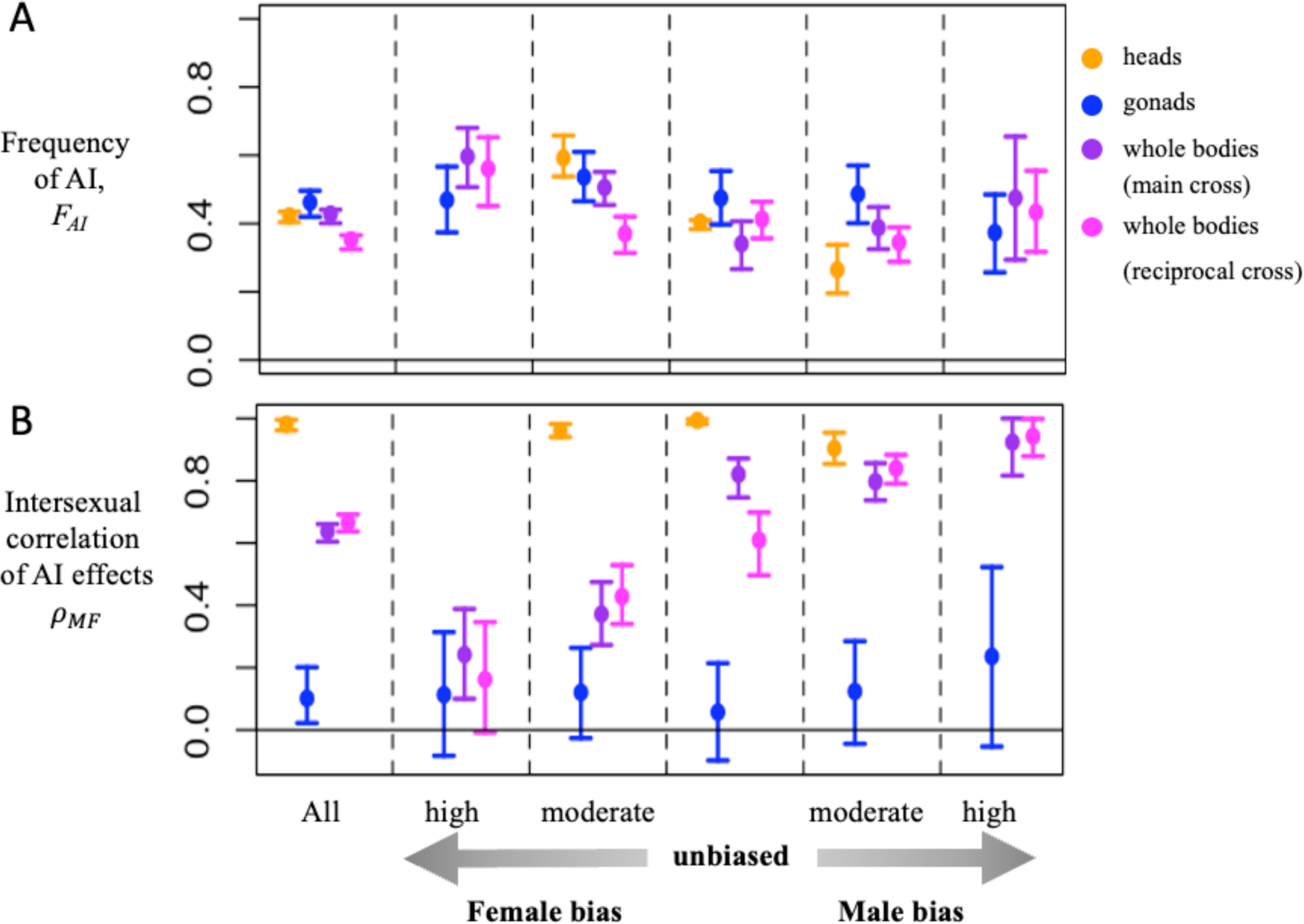
Model estimates of (A) the frequency of AI, *F_AI_*, and (B) the intersexual correlation of AI effects, *ρ_MF_*. The leftmost section in each panel shows results irrespective of sex-bias; the remaining sections show results stratified by sex bias: highly female-biased (-∞ < log2FC ≤ −2), moderately female-biased (−2 < log2FC ≤ −0.5), unbiased (−0.5 ≤ log2FC ≤ 0.5), moderately male-biased (0.5 < log2FC ≤ 2), highly male-biased (2 < log2FC < ∞). No analysis was performed for high sex-bias categories for heads as there were too few genes. Error bars represent 95% high density interval of the posterior. See Table S11 for other model parameters.

The intersexual correlation of AI effects is high in heads across all sex bias categories (*ρ_MF_* > 0.9; Fig. 1B) whereas it is much lower in gonads (*ρ_MF_* < 0.25). In the whole body samples, there is a striking change in *ρ_MF_* across sex-bias categories, going from low values for female-biased genes to high values for male-biased ones. (Applying the other model parameterization to these data, a qualitatively similar compatible pattern is manifest in that SD-AI is common among female-biased genes and much rarer for male-biased ones; Table S10).

### Sex-dependent reversals of allelic imbalance

Instances of sex-dependent allelic imbalance may include cases where the direction of imbalance is reversed between the sexes (i.e., male expression of the DGRP-177 allele is >50%, while female expression is <50%). Cases of sex-reversed AI are of special interest but are expected to be rare as this scenario presumably involves more complex regulation. While the Bayesian models in the previous sections are designed to make inferences about the distribution using information from a large number of genes, they are not well-suited to clearly identify individual genes exhibiting sex-reversed AI. To detect cases of sex-reversed AI, we analyzed genes individually, requiring significant AI effects in both sexes but of opposite direction. We examined whole body samples, combining data from both main and reciprocal crosses to maximize power. Of the 3796 genes tested in this fashion, 176 (4.6%) genes showed significantly sex-reversed AI. Using relatively conservative assumptions, we expect to find ∼26 cases by chance (i.e., false positives; see Supplementary Material).

We further examined these 176 genes using the head and gonad data to see if similar patterns can be observed in narrower tissue samples, i.e., gonads or heads. In the gonads, 155 of 176 genes had sufficient data to be examined for AI, whereas 160 genes could be examined in the heads. Of the 155 genes examined in the gonads, the majority (109) had sex-reversed point estimates of AI in the same direction as observed with the whole bodies whereas only 5 showed sex-reversed point estimates in the opposite direction relative to that found in the whole bodies. Thus, for the putative set of genes with sex-reversed AI, there exists a fair degree of concordance in the number and direction of sex-reversed AI between the gonads and whole bodies. In contrast, in heads, only 35 out of the 160 genes tested had sex-reversed point estimates of AI, and roughly half (only 17) of these were in the same direction as in the whole bodies, with the remaining half (18) in the opposite direction relative to whole bodies, i.e., not different from that expected by chance.

### Tissue-dependent patterns of allelic imbalance

In our previous analyses, we noted marked differences in the frequency of AI between heads and gonads. Though the analyses used different sets of genes for each tissue, the results suggest that AI may differ between tissues. To investigate tissue differences in AI more formally, we applied to the Bayesian model to estimate the joint distribution of AI effects in heads and gonads, separately for each sex (Table 3). In both sexes, we infer that ∼40% of genes exhibit AI and > 80% of these have effects that differ between gonads and heads. Using the alternative Bayesian model to characterize the distribution in terms of the between-tissue correlation in effects (Table S13), we find this correlation is quite low (*ρ_head-gonad_* ≈ 0.25) and that the average magnitude of effect sizes is larger in gonads than heads (gonads ≈ 0.09; heads ≈ 0.05).

**Table 3.** Parameter estimates [with 95% high posterior density interval] for the joint distribution of AI effects across heads and gonads.

| | Frequency<br>of genes<br>with AI,<br>$F_{AI}$ | Average<br>magnitude of<br>tissue-averaged AI<sup>1</sup>,<br>$\sigma_{AI}$ | Frequency of<br>genes with Tissue-Dependent AI<br>among genes with<br>AI,<br>$F_{TD-AI}$ | Average<br>magnitude of<br>tissue<br>difference in<br>AI<sup>2</sup>,<br>$\sigma_{TD-AI}$ | Overdispersion<br>parameter,<br>$\nu$ |
| --- | --- | --- | --- | --- | --- |
| Female | 0.37<br>[0.33, 0.41] | 0.053<br>[0.049, 0.057] | 0.94<br>[0.88, 1] | 0.094<br>[0.086, 0.101] | 0.0039<br>[0.0036, 0.0042] |
| Male | 0.44<br>[0.41, 0.47] | 0.050<br>[0.048, 0.053] | 0.84<br>[0.78, 0.91] | 0.099<br>[0.094, 0.105] | 0.0012<br>[0.001, 0.0014] |

<sup>1</sup> $\sigma_{AI}$ is the standard deviation of sex-averaged effects, which is modelled as approximately normally distributed with mean 0.
<sup>2</sup> $\sigma_{TD-AI}$ is the standard deviation of tissue differences in AI effects, which is modelled as approximately normally distributed with mean 0.

Analogous to our earlier test for sex-reversed AI, we tested for tissue-reversed AI, separately in each sex. We find 33 instances of possible tissue-reversed AI between gonads and heads in females, and 94 such instances in males. We estimate that 11 of these genes are likely false positives in females and 9 in males (see Supplementary Material). The much higher number of putative tissue-reversed AI in males may be a result of tissue differences in AI being larger in males, leading to easier detection of reversals in AI.

## Discussion

This study focuses on *cis*-regulatory variation in relation to sexual dimorphism in *D. melanogaster*. We utilise a divergent cross between North American (DGRP-177) and South African (SP159N) genotypes to maximize the number of polymorphic sites in the F_1_, thus increasing our power to detect allele-specific expression. To some extent, the patterns of allelic imbalance observed here may reflect inter-population differences in expression or may be idiosyncratic to the specific lines used here. As with any study of genetic variation, our results will be influenced by the interplay between the capacity of mutation to generate regulatory variation and selection to filter it. Our goal was to explore sex-specific patterns of *cis*-regulatory variation but our study was not designed to assess the relative importance of different evolutionary factors (e.g., mutation-selection-drift balance, balancing selection within populations, divergent selection among populations) in causing these patterns.

When analyzing individual genes, allelic imbalance was detected in 30-45% of genes analyzed (Table 1). These values are considerably higher than the values (6-17%) reported by Puixeu et al. (2023). The greater divergence of haplotypes used in our study may have contributed to this difference; we crossed fly lines from different continents whereas they crossed lines from within a single population. However, experimental and statistical differences are likely major contributors to this discrepancy. Rather than relying on the proportion of genes that reach significance in individual gene tests which depend heavily on statistical power, we also developed a Bayesian approach to directly estimate the frequency of genes with AI, finding that the frequency is quite high (38-48%, Table 2).

In heads and whole body samples, there is an enrichment of *cis*-regulatory variants in female-biased genes. Osada et al. (2017) and Puixeu et al. (2023) also examined *cis*-regulatory variation in relation to sex bias. Some of the patterns reported in those two studies appear similar while others appear different from ours or from each other. However, it is not possible to directly compare those studies to our own or to each other because of differences in methodology use to investigate these patterns. An advantage of our approach is that patterns can be investigated with respect to biologically interpretable properties such as the frequency of AI and average effect sizes.

The higher frequency of AI among female-biased genes that we detected may suggest that these genes are more frequently under balancing selection or tend to experience weaker purifying selection. Sexually antagonistic selection is one possible reason genes could experience balancing selection or weaker purifying selection. Cheng and Kirkpatrick (2016) suggested that genes with intermediate levels of sex-biased expression—both male-and female-biased—would be most likely to experience sexually antagonistic selection. As we do not observe an elevated frequency of AI for genes with intermediate levels of sex-bias, our results would seem to indicate that either that sexually antagonistic selection does not drive the pattern we observe or that the sexually antagonistic selection is not distributed across the genome as Cheng and Kirkpatrick (2016) suggested. While it is most intuitive to think of sexually antagonistic selection causing balancing selection or weak purifying selection, it could cause strong purifying selection if selection in one sex is much stronger than the other. Thus, the suggestion of Cheng and Kirkpatrick (2016) is not at necessarily at odds with our results if, for example, sexually antagonistic selection results in weak purifying selection for intermediately female-biased genes but strong purifying selection for intermediately male-biased genes.

Purifying selection (regardless of whether it involves sexual antagonism) is believed to be the most common form of selection and genes with *cis*-regulatory variation have previously been shown to be subject to weaker purifying selection in *Capsella* (Steige et al. 2017). Strongly female-biased genes in *D. melanogaster* have somewhat elevated levels of non-synonymous diversity (Singh and Agrawal 2023; but see details within), suggesting they may tend to be under weaker purifying selection and providing a potential explanation for the increased frequency of AI among these genes. A non-selective alternative explanation for our observed pattern is mutational bias; female-biased genes could have greater scope for *cis*-regulatory mutations (e.g., if such genes tend to have larger or more complex regulatory regions). We cannot ascertain which of these possibilities explain the observed pattern.

In general, *cis* effects are small in magnitude, with a large majority of genes exhibiting allelic expression differences below 5-10%. This finding is general to all the tissues examined, though average effects were notably higher in gonads than heads (Table S13). Others have also found frequent, weak *cis*-regulatory effects in other species, including *Capsella* (Steige et al. 2017), mice (Crowley et al. 2015) and *Saccharomyces* (Zhang and Emerson 2019). However, there have been reports of strong *cis* effects in *D. melanogaster* (León-Novelo et al., 2018), interspecific *Drosophila* hybrids (Graze et al. 2012) and natural populations of flycatchers (Wang et al. 2017), though the detection of low-effect variants may be encumbered by the lack of statistical power in some of these studies. An advantage of the Bayesian model approach we used is that it should allow us to infer the average effect size without the bias (“winner’s curse”) from gene-by-gene analyses.

Using gene-by-gene analyses, we found numerous instances of sex-dependent *cis*-regulatory effects in all tissues, though these were much more common in gonads than heads (Table 1). The inference from the Bayesian analysis indicated that almost all (>94%) *cis*-regulatory variants affect the sexes differently in gonads. Puixeu et al. (2023) also found a substantially higher frequency genes with sex differences in *cis*-regulatory effects in gonads than heads. The distinct regulatory architecture of the gonads, manifest in the high frequency of *cis*-regulatory variants with sex-dependent effects, should allow expression in gonads to evolve relatively independently between the two sexes whenever there is selective pressure to do so. By contrast, in heads we estimated that only a small fraction (∼5%) of *cis*-regulatory effects have sex-dependent effects, restricting the potential for independent evolution of expression between the two sexes. It is unsurprising that the frequency of sex-dependent AI in whole bodies is intermediate between the estimates inferred for gonads and heads. Relatedly, the intersexual correlation of AI effects, *ρ_MF_*, in whole bodies is intermediate between gonads and heads. However, there is a striking pattern of *ρ_MF_* in relation to sex bias in whole bodies that is not apparent in either gonads or heads: *ρ_MF_* steadily increases from strongly female-biased to strongly male-biased gene categories. This pattern seems similar to a pattern observed for the intersexual genetic correlation (*r_MF_*) in whole body gene expression (Singh and Agrawal 2023) in which the average *r_MF_* increases from strongly female-biased to strongly male-biased gene categories. It should be noted *ρ_MF_* and *r_MF_* are not defined at the same “level”; *ρ_MF_* is a correlation in AI effects between sexes estimated from a set of genes whereas each gene has its own *r_MF_* value—the intersexual correlation in expression level—estimated among a set of genotypes. Nonetheless, one would expect that if *ρ_MF_* was low (high) for a class of genes, then the average *r_MF_* for that class of genes would also be low (high) if *cis* regulatory effects were a major source of the genetic variation in expression. It is unclear why this pattern in *ρ_MF_* occurs in whole bodies but not heads or gonads and it is also unknown if *r_MF_* is related to sex-bias in tissues other than whole bodies.

Reversals in the direction of AI between the sexes are the most extreme form of sex-dependent AI and we detect such reversals at 176 loci in whole body samples. Puixeu et al. (2023) also documented some examples of sex-reversed AI (though using less stringent criteria to assess false positives). These are intriguing because heterozygotes of each sex will predominantly express alternative alleles for these genes. If the coding regions of the associated alleles were under sexually antagonistic selection, this type of *cis*-regulatory variation provides a potential mechanism by which sex-specific dominance reversals could occur (Grieshop et al. 2024). Dominance reversals have been invoked in the theoretical literature to explain the maintenance of genetic variation (Curtsinger et al., 1994; Kidwell et al., 1977; reviewed in Grieshop et al. 2024), though it has been unclear whether dominance reversals would be biologically plausible. Recently, this idea has gathered increasing empirical support with the discovery of sex-specific dominance reversal for traits likely under sexually divergent selection in salmonids (Barson et al. 2015; Pearse et al. 2019) as well as signals of dominance reversals for fitness in seed beetles (Grieshop and Arnqvist 2018). Sex reversals in AI potentially provide a mechanism, but it remains to be seen whether such *cis*-regulatory effects underlie any cases of dominance reversals for fitness.

In this study, we focussed on sex-dependent aspects of *cis*-regulatory variation because of its implications for sexual dimorphism and sexual conflict. However, ‘sex’ is one of many dimensions in which the context for a gene’s expression can differ. *Cis*-regulatory effects can vary within other contexts, such as temperature (Chen et al. 2015; Li and Fay 2017), tissue (Pinter et al. 2015), or metabolic state (Shih and Fay 2021). Of the two dimensions we examined—sex and tissue—neither stood out as harbouring dramatically more variation than the other (i.e., sex-dependent AI was very common in gonads as was tissue-dependent AI in both sexes). We were motivated to look for sex-dependent AI, at least in part, because of the possibility that sexually antagonistic selection might help maintain such variants. However, the qualitatively similar frequency of tissue-dependent AI suggests that either sexual antagonism is playing little role in maintaining regulatory variation genome-wide or that antagonistic pleiotropy across tissues is similarly common. Regardless, our results add to the existing literature that *cis*-regulatory effects are often context dependent in magnitude and sometimes even in direction. Moreover, different dimensions of context can interact to affect allelic expression, e.g., we find a sizeable number of genes with tissue-by-sex interaction effects on AI. It is plausible that this thinking extends to other contexts (e.g., sex-dependent *cis*-regulatory effects may be sensitive to temperature).

Allele-specific expression analysis can be plagued by inaccurate mapping of reads to the reference. We attempted minimize the influence of mapping bias in several ways. We constructed genotype-specific references and ascertained their efficacy by competitively mapping parental genomic reads to the references. We tested F_1_ genomic data for significant deviation from the expected read count ratio of 1:1 and removed any such gene from further consideration. We also tested for apparent ‘parental effects’ on allelic expression, which may ensue from any residual mapping bias, and excluded any such genes. Among our retained genes, the percentage of reads assigned to the DGRP-177 allele are uncorrelated between genomic and transcriptomic data, whereas a positive correlation would be expected if mapping bias was a substantial problem. The correlation remains negligible even if we only consider those genes with significant AI in expression. These steps are designed to allay the effects of mapping bias on our inferences, with evidence suggesting they have been largely effective (Supplementary Material). Nonetheless, we cannot entirely rule out the possibility that mapping bias may persist in the dataset within a small number of retained genes. While we believe our major findings are robust, the results for individual genes would benefit from further experimental validation.

As we infer *cis*-regulatory polymorphisms from allelic imbalance (i.e., relative expression) in heterozygotes, an additional note of caution is needed with respect to total expression. If allele *A* is expressed more than *B* in an *A/B* heterozygote, it is reasonable to assume that total expression of this gene would be greater in *A/A* than *B/B* homozygotes. However, this may not always be the case owing to negative feedback loops in transcription. Further experimental testing—including both homozygotes—would be needed to evaluate how often observed AI causes the predicted effects on total expression, though this is extremely difficult to do at scale while controlling for genetic background effects.

In summary, our data indicate that *cis-*regulatory variants are common and their effects are often dependent on sex and tissue. The frequency of *cis-*regulatory variants varies non-randomly with respect to sex-biased gene expression as well as across tissues. Further work remains to be done to more thoroughly examine variants at individual loci as well as to understand the evolutionary forces shaping patterns of variation genome-wide.

## Author Contributions

PM, KG, TB performed dissections and RNA extractions. PM performed bioinformatic data processing. PM and AA did the data analysis. PM, KG, and AA did the writing of the manuscript. AA conceived the experiment.

## Data availability

Upon acceptance, RNAseq data will be submitted to the Sequence Read Archive (NIH) and analysis scripts will be made available via GitHub.

## Acknowledgements

We thank Ina Anreiter for help with dissections, Baharul Choudhury for help with RNA extractions, Santiago Sánchez-Ramírez for bioinformatics advice, and Stephen Wright for helpful discussion. We thank John Pool and Yuheng Huang (U Madison) for providing the SP159N line.

