## Supplementary_Material for "*Cis-*regulatory variation in relation to sex and sexual dimorphism in *Drosophila melanogaster*"

Table of Contents


1. Supplemental text
    a. Text S1. Using genomic data to detect and assess mapping bias.
    b. Text S2. Consideration of false positive rates for tests of sex-reversed allelic imbalance
    c. Text S3. Details of Bayesian estimation of joint distribution of AI effects across the sexes
2. Supplemental tables
    a. Table S1. Filtering SNPs based on VCF annotation fields.
    b. Table S2. Summary of competitive mapping results for the RNA-seq samples.
    c. Table S3. Correlation of allele frequencies between genomic and transcriptomic reads among genes excluded from main analyses due to evidence of mapping bias.
    d. Table S4. Correlation of allele frequencies between genomic and transcriptomic reads from male F1 samples.
    e. Table S5. Correlation of allele frequencies between genomic and transcriptomic reads from female F1 samples.
    f. Table S6. Correlation of sex differences in allele frequencies estimated from genomic and transcriptomic data.
    g. Table S7. Fraction of genes with the given percentage of correctly assigned reads using parental genomic DNA.
    h. Table S8. Frequency of genes with AI and SD-AI among genes with ≥ 99% accuracy of parental genomic DNA read assignment.
    i. Table S9. Frequency of genes with AI and SD-AI using an FDR criterion.
    j. Table S10. Bayesian estimation of parameters for joint distribution of AI effects across the sexes using model framework 1 ("frequency of SD-AI" model).
    k. Table S11. Bayesian estimation of parameters for joint distribution of AI effects across the sexes using model framework 2 ("intersexual correlation of AI effects" model).
    l. Table S12. Pairwise differences [with 95% high posterior density interval] in $F_{AI}$ between different categories of sex bias.
    m. Table S13.

3. Supplemental figures
    a. Figure S1. Assignment of F1 genomic DNA reads.
    b. Figure S2. Allele frequencies estimated from genomic vs. transcriptomic reads from male F1 samples.
    c. Figure S3. Allele frequencies estimated from genomic vs. transcriptomic reads from female F1 samples.
    d. Figure S4. Sex differences in allele frequency estimated from genomic vs. transcriptomic reads from F1 samples.
    e. Figure S5. Distribution of the estimated frequency of DGRP-177 RNA-seq reads from whole body F1 samples, stratified with respect to parental genomic DNA read assignment accuracy.
    f. Figure S6. Distribution of the estimated frequency of DGRP-177 RNA-seq reads from whole body F1 samples using only genes with significant AI, stratified with respect to parental genomic DNA read assignment accuracy.

### Supplemental Text S1: Using genomic data to detect and assess mapping bias

#### *Using F1 genomic to identify and exclude genes with mapping bias*

Bias in mapping can lead to errors in estimation of allele-specific expression (Degner et al. 2009). Mapping bias can stem from multiple technical reasons, including sequence differences between the reads and reference genome, missing or false SNP calls and peculiarities in the alignment algorithm (León-Novelo et al. 2014). DNA controls were used to assess mapping bias in genes, and filter problematic genes from analysis. If mapping bias exists in a particular gene, paternal/maternal proportion of DNA read counts from $F_1$ heterozygotes will deviate from 50%. Our competitive mapping procedure was applied to genomic reads of $F_1$ heterozygotes (i.e., DNA controls). We aligned DNA to the genotype-specific references using BWA (Li and Durbin 2009).

For each gene with a total read count of >30, we calculated the binomial probability of the observed proportion of allele-specific read counts under the null expectation of 50% of each parental allele. (Genes with a total read count ≤ 30 in either male or female genomic data set were excluded from all analyses.) A gene was considered "suspicious" (i.e., potentially subject to mapping bias) if the observed proportion of parental reads fell in either 5% tail of the binomial distribution. We separately evaluated three genomic data sets: male genomic data, female genomic data, and combined data from both sexes. Using this criterion, we expect 10% of the genes in each data set to be identified as "suspicious" by chance (i.e., false positives). In each data set, we identify more than 10% of genes tested as suspicious (males: 12%; females: 13%; combined data: 14%). Genes that were identified as suspicious in any of these three data sets (3102 out of 12045 genes = 25%) were excluded from further analysis. Most genes had allelic coverage close to the expected 1:1 ratio (Figure S1). We also excluded genes where the estimated DGRP-177 coverage deviated from the expected frequency of DGRP-177 of 0.5 by more than 0.1, in either males or females. An additional 197 genes were excluded as a consequence of this criterion.

Despite these filters that aim to address mapping bias, it is possible that mapping bias still occurs for some of the retained genes. The list of genes identified with AI should be regarded only as candidates for genes with AI, rather than definitively being such. As an additional check, we examined the correlation between proportion of DGRP-177 in genomic and transcriptomic reads. A positive correlation is expected if mapping bias is falsely being interpreted as allelic imbalance.

For illustrative purposes, we first examine this correlation using the genes that were excluded from our main analyses because they had some evidence of mapping bias (though many of these excluded genes may be 'false positives', i.e., not truly have mapping bias). Specifically, we examine the correlation between the proportion of DGRP-177 in RNA-seq, as estimated from the quasibinomial model, and proportion of DGRP-177 in the genomic data. As expected with mapping bias, there are substantial positive correlations in both sexes and most tissues (Table S3). These correlations appear to be largely driven by the genes that would have met the statistical significance criteria for evidence of AI had they been included in the main analysis.  For reasons that are unclear, the genes that would have met the criteria for evidence of AI in the reciprocal cross have significant negative correlations.

When we instead consider the genes that were retained for the main analyses represent in Table 1, the correlations are near zero correlations in all tissues (Figures S2 & S3; Tables S4 & S5). These correlations are near zero both when including all genes as well as when considering only genes with evidence of AI. The sharp reduction in correlations post-filtering suggests that mapping bias is unlikely to be a major problem across the data set, but this does not preclude mapping bias affecting a relatively small set of genes.

As a major focus of our work is in sex differences in AI, we wanted to investigate whether sex differences in mapping bias could be a cause of a mis-inference of sex differences in AI. To do so, we examined the correlation of the sex difference in allele frequency in transcriptomic vs. genomic data. Mapping bias should cause a positive correlation. We found these correlations to be near zero (Figure S4; Table S6).

*Using parental genomic to assess the accuracy of competitive mapping*

To gauge if the genotype-specific references were sufficiently representative of the DGRP-177 and SP159N genotypes, we competitively mapped the parental (rather than F1) genomic reads to the two genotype-specific references. It is expected that few DGRP-177 reads should be competitively mapped to the SP159N reference, and vice-versa. For all genes with a coverage of ≥ 30, more than 99% genes had at least 90% reads mapped to the correct genotype-specific reference, while ~79% genes had at least 99% reads mapped correctly (Table S7). The rate of mis-mapping remained largely unchanged even as we filtered genes with apparent mapping bias or parental effects in allelic expression.

Nearly 20% of genes appear to have >1% reads which are mis-mapped, which may be a potential cause for concern. Such imperfect mapping could lead to false inferences of AI resulting from the mismapped reads. To assess whether the reported patterns of AI are affected by mismapped reads, we tested for AI in a set of genes that excluded any case where mismapping was >1%. We re-created the summary shown in Table 1 using this subset and is shown in Table S8. The similarity between Tables 1 and S8 indicates that the inclusion of genes where parental genomic read assignment was less accurate does not drastically skew the results.

To further assess any potential effects of mismapping, we calculated percentage of DGRP-177 reads for each gene using terms from the quasibinomial model performed on RNA-seq data from whole bodies. If mismapping is not a major issue, the proportion of DGRP-177 reads should lie in the vicinity of 0.5 for all genes. If substantial mapping issues persist in our data, we expect an upward bias (i.e., fraction of DGRP-177 in expression data to be > 0.5) for the set of genes where the DGRP-177 genomic data were assigned near-perfectly to the appropriate genome (>99% correctly-assigned reads) but the SP159N genomic data were not. Conversely, we expect a downward bias (i.e., fraction of DGRP-177 in expression data to be < 0.5) for the set of genes where the SP159N genomic data were assigned near-perfectly (>99% correctly-assigned reads) but the DGRP-177 genomic data were not. We observed that the proportion of DGRP-177 reads in the expression falls near 0.5 on average, regardless of whether or not there is a difference in how perfectly the two parental genomic data are assigned (Figure S6). This pattern also persists on examination of only genes with significant AI (Figure S7), though the pattern is a bit noisier due to the smaller number of genes.

Supplemental Text S2: Consideration of false positive rates for tests of sex-reversed allelic imbalance

Our criteria for sex-reversed AI (SRAI) are: a) significant AI effect using female data alone ($p < 0.05$), b) significant AI effect using male data alone ($p < 0.05$) and c) opposite directions of AI between the sexes. For the first two criteria, the probabilities of a false positive ($P_1$ and $P_2$, respectively) are 0.05, and the probability of AI being in opposite directions in the sexes ($P_3$) is 0.5. The expected number of false positives depends on the null hypothesis but, as discussed below, there is no simple obvious choice for this null hypothesis. We consider three different null hypotheses. None of these are likely to be correct. Our approach is to calculate the expected number of false positives under each of these hypotheses and then, conservatively, use the largest of these. The three null hypotheses we consider are as follows. Null hypothesis A: no AI in either sex (this might be considered the 'standard' null hypothesis); null hypothesis B: there exists AI in males but not females; and null hypothesis C: there exists AI in females, but not males. Null hypotheses B and C are conservative in the sense that they are *a priori* expected to yield high estimates for the number of false positives.

Under null hypothesis A, the probability of a false positive of SRAI is $P_A = P_1 \times P_2 \times P_3 = 0.00125$. The expected number of false positives is

$$E_A = 0.00125 \times n_T = 0.00125 \times 3796 = 4.75 \text{ genes},$$

where $n_T$ = total number of genes tested for AI.

Because we find numerous genes with AI, it seems unlikely that null hypothesis A is a good basis for assessing the expected number of false positives; it is too liberal and will underestimate the number of false positives. Null hypothesis B is much more conservative—arguably, too conservative—as it assumes the most extreme sex difference short of an actual sex reversal; thus, patterns suggestive of SRAI arise much more easily under null hypothesis B than A.

The probability of a false positive under null hypothesis B is $P_B = P_1 \times P_3 = 0.025$. The expected number of false positives is

$$E_B = 0.025 \times n_F = 0.025 \times 1805 = 45.125 \text{ genes},$$

where $n_F$ is the number of genes with significant AI in females.

Null hypothesis C is analogous to B. The probability of a false positive is $P_C = P_2 \times P_3 = 0.025$. The expected number of false positives is

$$E_C = 0.025 \times n_M = 0.025 \times 1813 = 45.325 \text{ genes}$$

where $n_M$ is the number of genes with significant AI in females.

The above estimates for number of false positives are likely to be too high because null hypotheses B and C assert that every gene with significant AI in one sex denotes a lack of AI in the other sex. It would be more realistic to assume that only a fraction *g* of the genes with AI in one sex represent cases where it is absent in the other. (Genes where there is truly AI in the *same* direction in both sexes are very unlikely to be sources of false positives for SRAI.)

Following this logic, the expected numbers of false positives under null hypotheses B and C are $E_B = 0.025 \times n_F \times g_F$ and $E_C = 0.025 \times n_M \times g_M$, where $g_F$ and $g_M$ represent the fraction of genes likely to give rise to false positives for females and males, respectively. We estimate this fraction for females as $g_F = 1 - f_F$ where $f_F$ = (number of genes with a significant sex effect on AI in analysis using both sexes as well significant AI in analysis of females alone) / (number of genes with significant AI in analysis of females alone). This should be a reasonably conservative estimate of $g_F$ (i.e., we are more likely to overestimate $g_F$ than underestimate it because of our limited power to detect sex-dependent AI). $g_M$ and $f_M$ are defined analogously. We have $f_F = 824/1805$, and $f_M = 823/1813$. Substituting these values in the expressions above, we have:

$$E_B = 1805 \text{ x } (1 - 824/1805) \text{ x } 0.05 \text{ x } 0.5 = 24.525 \text{ genes}$$

and

$$E_C = 1813 \text{ x } (1 - 823/1813) \text{ x } 0.05 \text{ x } 0.5 = 24.75 \text{ genes}$$

Taking the larger of the two expected values and rounding up, we expect 25 false positive cases of sex-reversed allelic imbalance.

The expected numbers of false positives for tissue-reversed AI were estimated in a similar fashion: $E_B = 0.025 \text{ x } n_G \text{ x } g_G$ and $E_C = 0.025 \text{ x } n_H \text{ x } g_{H,}$ where $n_G$, $g_G$, $n_H$ and $g_H$ denote terms congruent to that for the sex-specific AI analysis, but for gonads and heads. The expected numbers of false positives in the analysis for females are:

$$E_B = 0.025 \text{ x } n_H \text{ x } g_H = 703 \text{ x } (1 - 274/703) \text{ x } 0.05 \text{ x } 0.5 = 10.725 \text{ genes}$$

and

$$E_C = 0.025 \text{ x } n_G \text{ x } g_G = 380 \text{ x } (1 - 216/380) \text{ x } 0.05 \text{ x } 0.5 = 4.10 \text{ genes}$$

Likewise, for males:

$$E_b = 0.025 \text{ x } n_H \text{ x } g_H = 761 \text{ x } (1 - 413/761) \text{ x } 0.05 \text{ x } 0.5 = 8.7 \text{ genes}$$

and

$$E_c = 0.025 \text{ x } n_G \text{ x } g_G = 1104 \text{ x } (1 - 752/1104) \text{ x } 0.05 \text{ x } 0.5 = 8.8 \text{ genes}$$

Taking the larger of the two expected values for each sex and rounding, we use 11 and 9 as the expected number of false positive cases of tissue-reversed allelic imbalance in females and males, respectively.

Supplement Text S3. Details of Bayesian estimation of joint distribution of AI effects across the sexes

*Model framework 1*

Let $f_{S,i}$ be the proportion of expression from the DGRP-177 allele in sex $s$ for gene $i$. The sex-averaged AI for gene $i$ is $a_i = \frac{f_{F,i}+f_{M,i}}{2} - \frac{1}{2}$. The sex difference in AI for gene $i$ is $d_i = f_{F,i} - f_{M,i}$.

Let $F_{AI}$ be the faction of genes that have a non-zero value of sex-averaged AI. For for this set of genes, we assume that values of $a$ are 'approximately' (see below) normally distributed with mean 0 (i.e., no genome-wide bias towards one strain's alleles) and standard deviation, $\sigma_{AI}$. Among those genes with non-zero sex-averaged AI, the fraction of genes with a sex difference in AI is given by $F_{SD\text{-}AI}$. For these genes, we assume values of $d$ are approximately normally distributed with mean 0 and standard deviation, $\sigma_{SD\text{-}AI}$. (Technically, the model assumes there are no genes with exactly equal but opposite AI between the sexes, i.e., where $a = 0$ but $d \neq 0$. Practically, such genes can be approximately accommodated in this model by small values of sex-averaged AI with non-zero sex differences in AI.) For practical reasons, we use discrete distributions (based on normal distributions) for both $a$ and $d$. Magnitudes of $a$ (i.e., $|a|$) come from the set $x_a$ = {0.01, 0.02, 0.04, 0.07, 0.1, 0.15, 0.2, 0.25, 0.3, 0.35, 0.4, 0.45}. The probability that $|a|$ is equal to the $j^{th}$ element of this set is

$$P(|a| = x_a[j]) = \frac{CDF_{0,\sigma_{AI}}\left[x_a[j]\right] - CDF_{0,\sigma_{AI}}\left[x_a[j-1]\right]}{\sum_{k=1}(CDF_{0,\sigma_{AI}}\left[x_a[k]\right] - CDF_{0,\sigma_{AI}}\left[x_a[k-1]\right])}$$

where $x_a[0] \equiv 0$ and $CDF_{0,\sigma_{AI}}[x]$ is the cumulative density function for a normal distribution with mean 0 and standard deviation $\sigma_{AI}$. Larger values of $\sigma_{AI}$ increase the probability mass associated with larger values of $|\alpha|$. With equal probability, $a = |a|$ or $a = -|a|$, i.e., the distribution of $a$ is symmetric around 0.

Values for $d$ are modelled similarly. Magnitudes of $d$ (i.e., $|d|$) come from the set $x_d$ = {0.02, 0.04, 0.06, 0.09, 0.12, 0.15, 0.2, 0.25, 0.3, 0.4, 0.5, 0.7}. The probability that $|d|$ is equal to the $j^{th}$ element of this set is

$$P(|d| = x_d[j]) = \frac{CDF_{0,\sigma_{SD-AI}}\left[x_d[j]\right] - CDF_{0,\sigma_{SD-AI}}\left[x_d[j-1]\right]}{\sum_{k=1}(CDF_{0,\sigma_{SD-AI}}\left[x_d[k]\right] - CDF_{0,\sigma_{SD-AI}}\left[x_d[k-1]\right])}$$

where $x_d[0] \equiv 0$ and $CDF_{0,\sigma_{SD-AI}}[x]$ is the cumulative density function for a normal distribution with mean 0 and standard deviation $\sigma_{SD-AI}$. With equal probability, $d = |d|$ or $d = -|d|$, i.e., the distribution of $d$ is symmetric around 0.

The values of $a$ and $d$ are used to determine $f_F$ and $f_M$: $f_F = 0.5 + a + d/2$ and $f_M = 0.5 + a - d/2$. Some combinations of values for $a$ and $d$ result in either $f_F$ and $f_M$ outside the permissible range, i.e., [0, 1]; in these cases, the value is set to the boundary value. For realistic values of the relevant parameters, $\sigma_{AI}$ and $\sigma_{SD-AI}$, there is very little probability mass associated with these extremes.

A summary of the joint distribution of $f_F$ and $f_M$ is given in the table below.

| Comment | $\{f_F, f_M\}$ | $P_{f,f}(f_F, f_M)$ |
|---|---|---|
| No AI | {0.5, 0.5} | $1 - F_{AI}$ |
| AI but no SD-AI | $\{0.5 + a, 0.5 + a\}$ | $F_{AI}(1 - F_{SD-AI})\ P(a)$ |
| AI with SD-AI | $\{0.5 + a + d/2,$<br>$0.5 + a - d/2\}$ | $F_{AI}F_{SD-AI}\ P(a)P(d)$ |

The likelihood of the data is

$L[data|F_{AI}, F_{SD-AI}, \sigma_{AI}, \sigma_{SD-AI}, v] =$

$$\prod_{i=1}^{N_{genes}} \sum_{\{f_F, f_M\}} P_{f,f}(f_F, f_M)\left(\prod_{j=1}^{3} DBB[n_{1,i,F,j}, n_{2,i,F,j}, \alpha[f_F, v], \beta[f_F, v]]\right)\left(\prod_{k=1}^{3} DBB[n_{1,i,M,k}, n_{2,i,M,k}, \alpha[f_M, v], \beta[f_M, v]]\right)$$

The first product operator is over all genes being analyzed and the first summation is over all combinations of $f_F$ and $f_M$. (As described above, the joint distribution of $f_F$ and $f_M$ is determined by $F_{AI}, F_{SD-AI}, \sigma_{AI}$, and $\sigma_{SD-AI}$.) $n_{1,i,S,j}$ and $n_{2,i,S,j}$ are the numbers of reads of allele 1 and allele 2, respectively, for gene *i* from sample *j* of sex *S*. $DBB[x, y, \alpha, \beta]$ is the probability of observing *x* successes and *y* failures from a beta binomial distribution with parameters $\alpha$ and $\beta$. We express $\alpha$ and $\beta$ as functions: $\alpha[f, v] = f(\frac{1}{v} - 1)$ and $\beta[f, v] = (1 - f)(\frac{1}{v} - 1)$. The parameter $v$ captures overdispersion; when $v = 0$, *DBB* becomes equivalent to a binomial distribution with probability of success *f* but with $v > 0$ there is more variance among samples than expected under the binomial distribution.

*Model framework 2*

Let $F_{AI}$ be the faction of genes that have a non-zero value for AI (as in the first model framework). For such genes, let $a_{F,i}$ represent the deviation from zero imbalance in females, i.e., $a_{F,i} = f_{F,i} - \frac{1}{2}$; $a_{M,i}$ is the corresponding values in males. We assume values of $\{a_F, a_M\}$ are approximately bivariate normally distributed with mean 0 and with standard deviations $\sigma_{AI,F}$ and $\sigma_{AI,M}$ for females and males, respectively, and an intersexual correlation of $\rho_{MF}$. We use discretized approximation to the bivariate normal distribution. Magnitudes of both $a_F$ and $a_M$ (i.e., $|a_F|$ and $|a_M|$) come from the set $x_a$ = {0.01, 0.02, 0.04, 0.07, 0.1, 0.15, 0.2, 0.25, 0.3, 0.35, 0.4, 0.45}. The probability that $a_F = x_a[j]$ and $a_M = x_a[k]$ is proportional to

$$P(a_F = x_a[j], a_M = x_a[k]) = \frac{\int_{x_a[j-1]}^{x_a[j]} \int_{x_a[k-1]}^{x_a[k]} PDF_{0,\boldsymbol{V}}[x,y]\partial x \partial y}{T}$$

where $x_a[0] \equiv 0$ and $PDF_{\vec{0},\boldsymbol{V}}[x,y]$ is the probability density function for a bivariate normal distribution with mean {0, 0} and variance-covariance matrix

$$\boldsymbol{V} = \begin{pmatrix} \sigma_{AI,F}^2 & \rho_{MF}\sigma_{AI,F}\sigma_{AI,M} \\ \rho_{MF}\sigma_{AI,F}\sigma_{AI,M} & \sigma_{AI,M}^2 \end{pmatrix}$$

$T$ is a normalization factor such that the sum over all allowable values of $P(a_F, a_M)$ is 1. The equation above applies if both $a_F$ and $a_M$are positive because all values in $x_a$ are positive. By symmetry of the bivariate normal distribution, $P(a_F = -x_a[j], a_M = -x_a[k]) = P(a_F = x_a[j], a_M = x_a[k])$. If $a_F > 0$ and $a_M < 0$, then

$$P(a_F = x_a[j], a_M = -x_a[k]) = \frac{\int_{x_a[j-1]}^{x_a[j]} \int_{-x_a[k]}^{-x_a[k-1]} PDF_{0,\boldsymbol{V}}[x,y]\partial x \partial y}{T}$$

If $a_F > 0$ and $a_M < 0$, then

$$P(a_F = -x_a[j], a_M = x_a[k]) = \frac{\int_{-x_a[j]}^{-x_a[j-1]} \int_{x_a[k-1]}^{x_a[k]} PDF_{0,\boldsymbol{V}}[x,y]\partial x \partial y}{T}$$

A summary of the joint distribution of $f_F$ and $f_M$ for this model framework is given in the table below.

| Comment | $\{f_F, f_M\}$ | $P_{f,f}(f_F, f_M)$ |
|---|---|---|
| No AI | {0.5, 0.5} | $1 - F_{AI}$ |
| AI | {0.5 + $a_F$, 0.5 + $a_M$} | $F_{AI}\ P(a_F, a_M)$ |

The likelihood of the data is calculated as described above for the first model framework but calculating $P_{f,f}(f_F, f_M)$ as described in this section.

*Additional data filtering for Bayesian analysis*

Genes for which there is more data (i.e., more reads) will have a disproportionate effect on the inferred distribution of AI effects. This could be problematic if, for example, the AI properties of highly expressed genes are not representative of the overall distribution. To minimize this issue, we trimmed the data as follows, separately for each tissue type. First we excluded genes that were in the bottom quartile with respect to the total number of (assignable) reads summed across all three replicates for either sex. (This excludes a larger fraction of genes in gonads than heads because there is less overlap in which genes are in the bottom quartile for males versus females in gonads than heads. Nonetheless, there are still many more sex-biased genes in gonads than heads even after this process.) Then, separately for each sex, we down-sampled such that the total read count summed across all three replicates was the same for each gene. The down-sampling was performed to improve the balance in total count across the three replicates. For example, if the original total read count was 93, 120, 163 for the three replicates and the goal was to down-sample to a combined total of 300, only the latter two replicates would be down-sampled (e.g., read counts after down-sample: 93, 104, 103). (Though it is too computationally demanding to fully examine the effect of this data filtering, we did perform a cursory inspection. While there were some quantitative differences in parameter estimates between using the full data versus this trimmed data, the parameter estimates—and the striking differences between tissue types—were qualitatively similar between the full and filtered data sets.) For analyses performed on genes within a specified sex-bias category, the filtering described above was applied first (i.e., considering all genes irrespective of sex-bias), and then the remaining genes post-filtering were sorted into their respective sex-bias categories. Analyses were performed on these subsetted genes.

The calculation of the likelihood for any one parameter combination is computationally slow. For this reason, we first used the Nelder-Mead algorthim in *optim*, from multiple different starting parameter values, to find a parameter combination in the vicinity of the maximum likelihood parameter combination and then used this as the starting value for MCMC approximation of the posterior distribution obtained from R package *BayesianTools* (Hartig et al. 2023). For the first model framework, we used (independent) uniform priors for each of the five parameters on the following intervals: $F_{AI} \in [0,1], F_{SD-AI} \in [0,1]$, $\sigma_{AI} \in [0,0.5]$, $\sigma_{SD-AI} \in [0,0.5]$, and $\upsilon \in [0,0.1]$. For the second model framework, we used (independent) uniform priors for each of the five parameters on the following intervals: $F_{AI} \in [0,1]$, $\sigma_{AI,F} \in [0,0.5], \sigma_{AI,M} \in [0,0.5]$, $\rho_{MF} \in [-1,1]$, and $\upsilon \in [0,0.1]$. For each tissue data set, we ran six MCMC chains. Each chain had 10000 iterations. After removing the first 500 samples from each chain and then applying a thinning interval of 5, the six chains were combined to construct the final chain. The marginal posterior distributions for each parameter are drastically different than the corresponding priors, indicating that the priors cause negligible bias in the posteriors.

**Table S1.** Filtering SNPs based on VCF annotation fields. We followed GATK's generic recommendations for hard-filtering short variants. SNPs that failed to meet any of the thresholds for VCF annotation fields were left out from being updated into the genotype-specific references. The sites corresponding to these filtered SNPs were left unchanged in the reference genomes and not masked (i.e., both genotype-specific references retained the original *D. melanogaster* Release 6 reference genome sequence at such sites).

| Annotation field | Filtering criterion |
|---|---|
| QualByDepth (QD) | >2 |
| Fisher Strand (FS) | >60 |
| RMS Mapping Quality (MQ) | >40 |
| Mapping Quality Rank Sum Test (MQRankSumTest) | <-12.5 |
| Read Position Rank SumTest (ReadPosRankSum) | <-8 |

**Table S2**. Summary of competitive mapping results for the RNA-seq samples. The table shows the proportion of assignable reads retained for each sample in the study. A read is retained if it can be assigned to either parental reference genome based on the corresponding values of the Alignment Score (AS) and/or number of mismatches (nM). As evident from the table, more than half of RNA-seq reads are ambiguous (i.e., had identical values of AS and nM when aligned to each reference), leading them to be unassigned.

| Sample | Total number of mapped reads | Number of assignable reads (DGRP-177 + SP159N) | % assignable reads |
|---|---|---|---|
| Female heads 1 | 97891874 | 42078498 | 43% |
| Female heads 2 | 191809824 | 80257450 | 42% |
| Female heads 3 | 174137048 | 74833400 | 43% |
| Male head 1 | 86403742 | 36931962 | 43% |
| Male head 2 | 67410878 | 28278228 | 42% |
| Male head 3 | 73184204 | 31223677 | 43% |
| Male whole bodies (reciprocal) 1 | 64162128 | 26101256 | 41% |
| Male whole bodies (reciprocal) 2 | 103771934 | 44046474 | 42% |
| Male whole bodies (reciprocal) 3 | 198399718 | 81826628 | 41% |
| Male whole bodies 1 | 107285194 | 45376756 | 42% |
| Male whole bodies 2 | 63607856 | 26738694 | 42% |
| Male whole bodies 3 | 61017004 | 25771904 | 42% |
| Female whole bodies (reciprocal) 1 | 171499724 | 66846478 | 39% |
| Female whole bodies (reciprocal) 2 | 168366246 | 48442330 | 29% |
| Female whole bodies (reciprocal) 3 | 161579810 | 59316712 | 37% |
| Female whole bodies 1 | 193008604 | 71283018 | 37% |
| Female whole bodies 2 | 87117482 | 31008324 | 36% |
| Female whole bodies 3 | 74792732 | 18388580 | 25% |

| | | | |
|---|---|---|---|
| Testes 1 | 54620538 | 25947704 | 47% |
| Testes 2 | 60525228 | 29206388 | 48% |
| Testes 3 | 108260568 | 51577232 | 48% |
| Ovaries 1 | 82143190 | 16763652 | 20% |
| Ovaries 2 | 89799596 | 16621676 | 18% |
| Ovaries 3 | 74198336 | 24360660 | 33% |

**Table S3**. Correlation of allele frequencies between genomic and transcriptomic reads among genes excluded from main analyses due to evidence of mapping bias. Significant correlations are denoted in bold; Pearson's correlation tests, $p < 0.05$. Numbers in parentheses represent the number of genes for the given correlation.

| Sample | Genes that would have met the criterion for AI | | Genes that would *not* have met the criterion for AI | | All genes | |
|---|---|---|---|---|---|---|
| | Females | Males | Females | Males | Females | Males |
| Gonads | **0.356** (787) | **0.076** (787) | **0.079** (1075) | **0.060** (1075) | **0.249** (1862) | **0.079** (1862) |
| Heads | **0.238** (1577) | **0.052** (1577) | 0.035 (1265) | -0.036 (1265) | **0.182** (2842) | **0.050** (2842) |
| Whole bodies (main) | **0.247** (1494) | **0.053** (1494) | 0.053 (1270) | -0.05 (1270) | **0.192** (2764) | **0.051** (2764) |
| Whole bodies (reciprocal) | **-0.151** (1565) | -0.04 (1563) | **-0.008** (1375) | 0.003 (1375) | **-0.119** (2938) | **-0.044** (2938) |

**Table S4.** Correlation of allele frequencies between genomic and transcriptomic reads from male F1. These analyses are using the set of genes post-filtering, i.e., the set of genes for the results shown in Table 1. The value shown in bold has $0.01 < p < 0.05$ in a Pearson's correlation test. Though this one correlation is significant, the estimated magnitude is small and this correlation would not be significant when correcting for multiple tests. Numbers in parentheses represent the number of genes for the given correlation.

| Sample | Genes with AI | Genes with no AI | All genes |
|---|---|---|---|
| Gonads | 0.009 (805) | -0.019 (1742) | -0.003 (2547) |
| Heads | -0.012 (1830) | -0.013 (2333) | -0.011 (2063) |
| Whole Bodies (main) | 0.047 (1325) | 0.007 (2303) | 0.027 (3628) |
| Whole Bodies (reciprocal) | **0.058** (1392) | -0.037 (2509) | 0.017 (3901) |

**Table S5.** Correlation of allele frequencies between genomic and transcriptomic reads from female F1 samples. These analyses are using the set of genes post-filtering, i.e., the set of genes for the results shown in Table 1. None of these correlations are significantly different than zero ($p > 0.05$; Pearson's correlation test). Numbers in parentheses represent the number of genes for the given correlation.

| Sample | Genes with AI | Genes with no AI | All genes |
|---|---|---|---|
| Gonads | -0.009 (805) | 0.003 (1742) | -0.002 (2547) |
| Heads | -0.012 (1830) | 0.006 (2333) | -0.006 (2063) |
| Whole Bodies (main) | 0.017 (1325) | -0.013 (2303) | 0.005 (3628) |
| Whole Bodies (reciprocal) | 0.040 (1392) | -0.029 (2509) | 0.011 (3901) |

**Table S6.** Correlation of sex differences in allele frequencies estimated from genomic and transcriptomic data. These analyses are using the set of genes post-filtering, i.e., the set of genes for the results shown in Table 1. None of these correlations are significantly greater than zero ($p > 0.05$; Pearson's correlation test). Numbers in parentheses represent the number of genes.

| Sample | Genes with AI, without SDAI | Genes without AI, with SDAI | Genes with AI and SDAI | Genes without AI or SDAI | All genes |
|---|---|---|---|---|---|
| Gonads | -0.024 (409) | -0.056 (249) | -0.003 (396) | 0.006 (1493) | -0.006 (2547) |
| Heads | -0.013 (1526) | -0.060 (106) | 0.079 (304) | 0.004 (2127) | 0.006 (4063) |
| Whole bodies (main) | 0.022 (871) | -0.057 (256) | 0.020 (452) | -0.017 (2047) | -0.004 (3636) |
| Whole bodies (reciprocal) | 0.021 (943) | -0.023 (265) | 0.048 (449) | -0.047* (2244) | -0.008 (3901) |

**Table S7.** Fraction of genes with the given percentage of correctly assigned reads using parental genomic DNA. Genomic DNA for each of the two parental genotypes was subject to the competitive mapping procedure.

| % correctly mapped reads | DGRP-177 | | SP159N | |
|---|---|---|---|---|
| | Fraction of genes, pre-filtering | Fraction of genes, post-filtering | Fraction of genes, pre-filtering | Fraction of genes, post-filtering |
| ≥ 90% | 0.998 | 0.999 | 0.996 | 0.997 |
| ≥ 95% | 0.993 | 0.995 | 0.989 | 0.991 |
| ≥ 99% | 0.795 | 0.797 | 0.785 | 0.793 |

**Table S8.** Frequency of genes with AI and SD-AI among genes with ≥ 99% accuracy of parental genomic DNA read assignment. Genes with AI and SD-AI as identified from generalized linear models on individual genes, as in Table 1.

| Tissue | Number of genes tested | % genes with AI | % genes with sex-dependent AI |
|---|---|---|---|
| gonads | 1691 | 32.6% | 26.4% |
| heads | 2660 | 46.2% | 10.2% |
| whole bodies (main) | 2403 | 38.0% | 21.1% |
| whole bodies (reciprocal) | 2582 | 36.2% | 18.6% |

**Table S9**. Frequency of genes with AI and SD-AI using an FDR criterion. This table is based on the set of genes analyzed with GLMs as in Table 1. Table 1 is based on defining genes as "significant" if the appropriate term has a *p*-value < 0.05. For this table, false discovery rate *q*-values were calculated using the base *R* function for Benjamini-Hochberg procedure and genes are defined as "significant" if the appropriate term has a *q*-value < 0.05.

| Tissue | Number of genes tested | % genes with AI | % genes with sex-dependent AI |
|---|---|---|---|
| gonads | 2547 | 19.1% | 12.8% |
| heads | 4063 | 35.9% | 2.7% |
| whole bodies (main) | 3628 | 26.3% | 9.1% |
| whole bodies (reciprocal) | 3901 | 25.8% | 7.3% |

**Table S10.** Bayesian estimation of parameters [with 95% high posterior density interval] for joint distribution of AI effects across the sexes using model framework 1 ("frequency of SD-AI" model).

| | $F_{AI}$[1] | $F_{SD\text{-}AI}$[2] | $\sigma_{AI}$[3] | $\sigma_{SD\text{-}AI}$[4] | $\upsilon$[5] |
|---|---|---|---|---|---|
| **heads** | | | | | |
| All genes $n$ = 2910 | 0.42 [0.40, 0.45] | 0.05 [0.01, 0.08] | 0.057 [0.054, 0.06] | 0.074 [0.051, 0.106] | 0.001 [0.0008, 0.0011] |
| moderate FB $n$ = 484 | 0.60 [0.52, 0.67] | 0.12 [0.04, 0.22] | 0.058 [0.052, 0.063] | 0.062 [0.039, 0.091] | 0.0009 [0.0006, 0.0012] |
| UB $n$ =1877 | 0.40 [0.37, 0.44] | 0.05 [0.01, 0.1] | 0.057 [0.054, 0.061] | 0.055 [0.032, 0.078] | 0.001 [0.0008, 0.0011] |
| moderate MB $n$ = 535 | 0.30 [0.23, 0.38] | 0.02 [0, 0.05] | 0.055 [0.048, 0.062] | 0.253 [0.142, 0.384] | 0.0012 [0.0009, 0.0015] |
| **gonads** | | | | | |
| All genes $n$ = 1550 | 0.48 [0.45, 0.52] | 0.94 [0.9, 1] | 0.053 [0.05, 0.056] | 0.105 [0.099, 0.114] | 0.0056 [0.0053, 0.006] |
| high FB $n$ = 246 | 0.49 [0.39, 0.58] | 0.92 [0.79, 1] | 0.066 [0.056, 0.077] | 0.132 [0.111, 0.155] | 0.0086 [0.0075, 0.0098] |
| moderate FB $n$ =421 | 0.57 [0.49, 0.65] | 0.86 [0.72, 1] | 0.05 [0.044, 0.057] | 0.104 [0.091, 0.119] | 0.0062 [0.0056, 0.007] |
| UB $n$ = 370 | 0.49 [0.41, 0.57] | 0.95 [0.87, 1] | 0.054 [0.047, 0.061] | 0.112 [0.098, 0.127] | 0.0055 [0.0048, 0.0063] |
| moderate MB $n$ =346 | 0.51 [0.41, 0.59] | 0.88 [0.75, 1] | 0.042 [0.036, 0.048] | 0.086 [0.073, 0.101] | 0.0041 [0.0035, 0.0048] |
| high MB $n$ = 167 | 0.40 [0.27, 0.52] | 0.80 [0.59, 1] | 0.045 [0.035, 0.055] | 0.085 [0.063, 0.108] | 0.0026 [0.0018, 0.0034] |

continued

**Table S10** continued.

| | $F_{AI}$[1] | $F_{SD\text{-}AI}$[2] | $\sigma_{AI}$[3] | $\sigma_{SD\text{-}AI}$[4] | $\upsilon$[5] |
|---|---|---|---|---|---|
| **whole body (main cross)** | | | | | |
| All genes *n* = 2170 | 0.35 [0.29, 0.41] | 0.72 [0.55, 0.88] | 0.068 [0.06, 0.077] | 0.054 [0.044, 0.064] | 0.0019 [0.0016, 0.0022] |
| high FB *n* = 313 | 0.64 [0.54, 0.81] | 0.81 [0.52, 1] | 0.028 [0.024, 0.032] | 0.054 [0.044, 0.065] | 0.0012 [0.001, 0.0016] |
| moderate FB *n* = 888 | 0.53 [0.46, 0.58] | 0.76 [0.66, 0.85] | 0.035 [0.032, 0.038] | 0.061 [0.056, 0.067] | 0.0014 [0.0012, 0.0016] |
| UB *n* = 509 | 0.35 [0.29, 0.41] | 0.72 [0.55, 0.88] | 0.068 [0.06, 0.077] | 0.054 [0.044, 0.064] | 0.0019 [0.0016, 0.0022] |
| moderate MB *n* = 389 | 0.43 [0.35, 0.50] | 0.38 [0.26, 0.48] | 0.067 [0.059, 0.077] | 0.078 [0.061, 0.098] | 0.0016 [0.0013, 0.002] |
| high MB *n* = 77 | 0.49 [0.33, 0.69] | 0.34 [0, 0.66] | 0.045 [0.032, 0.06] | 0.027 [0, 0.076] | 0.0012 [0.0006, 0.0019] |
| **whole body (reciprocal cross)** | | | | | |
| All genes *n* = 2386 | 0.38 [0.35, 0.4] | 0.66 [0.60, 0.71] | 0.055 [0.053, 0.058] | 0.065 [0.06, 0.070] | 0.0026 [0.0025, 0.0027] |
| high FB *n* = 272 | 0.61 [0.5, 0.75] | 0.74 [0.53, 0.98] | 0.024 [0.021, 0.029] | 0.055 [0.044, 0.069] | 0.0023 [0.0018, 0.0026] |
| moderate FB *n* = 858 | 0.38 [0.33, 0.43] | 0.82 [0.70, 0.94] | 0.045 [0.040, 0.049] | 0.068 [0.061, 0.077] | 0.0028 [0.0025, 0.0031] |
| UB *n* =572 | 0.42 [0.36, 0.48] | 0.76 [0.63, 0.89] | 0.042 [0.038, 0.047] | 0.051 [0.043, 0.06] | 0.002 [0.0018, 0.0023] |
| moderate MB *n* =530 | 0.37 [0.31, 0.43] | 0.35 [0.20, 0.49] | 0.085 [0.075, 0.095] | 0.087 [0.065, 0.110] | 0.0029 [0.0026, 0.0032] |
| high MB *n* =154 | 0.44 [0.33, 0.58] | 0.42 [0.04, 0.79] | 0.050 [0.039, 0.060] | 0.034 [0, 0.076] | 0.0025 [0.0019, 0.003] |

[1] $F_{AI}$ = frequency of genes with AI

[2] $F_{SD\text{-}AI}$ = frequency of genes with SD-AI among genes with AI

[3] $\sigma_{AI}$ = average magnitude of sex-averaged AI; $\sigma_{AI}$ is the standard deviation of sex-averaged effects, which is modelled as approximately normally distributed with mean 0

[4] $\sigma_{SD\text{-}AI}$ = average magnitude of sex-differences in AI effects for genes with SD-AI; $\sigma_{SD\text{-}AI}$ is the standard deviation of sex-differences in AI effects, which is modelled as approximately normally distributed with mean 0

[5] $\upsilon$ = overdispersion parameter

**Table S11**. Bayesian estimation of parameters [with 95% high posterior density interval] for joint distribution of AI effects across the sexes using model framework 2 ("intersexual correlation of AI effects" model).

| | $F_{AI}$[1] | $\sigma_{AI,F}$[2] | $\sigma_{AI,M}$[3] | $\rho_{MF}$[4] | $\upsilon$[5] |
|---|---|---|---|---|---|
| heads | | | | | |
| All genes *n* = 2910 | 0.42 [0.4, 0.43] | 0.058 [0.055, 0.061] | 0.058 [0.055, 0.061] | 0.98 [0.96, 1] | 0.001 [8e-04, 0.001] |
| moderate FB *n* = 484 | 0.59 [0.54, 0.66] | 0.062 [0.056, 0.067] | 0.059 [0.053, 0.063] | 0.96 [0.94, 0.98] | 8e-04 [5e-04, 0.0011] |
| UB *n* =1877 | 0.40 [0.38, 0.41] | 0.057 [0.056, 0.059] | 0.058 [0.058, 0.06] | 0.99 [0.98, 1] | 0.001 [8e-04, 0.0011] |
| moderate MB *n* = 535 | 0.26 [0.2, 0.34] | 0.06 [0.05, 0.07] | 0.058 [0.049, 0.066] | 0.9 [0.85, 0.95] | 0.0012 [9e-04, 0.0015] |
| gonads | | | | | |
| All genes *n* = 1550 | 0.46 [0.42, 0.5] | 0.076 [0.072, 0.081] | 0.074 [0.069, 0.079] | 0.1 [0.02, 0.2] | 0.0057 [0.0053, 0.0061] |
| high FB *n* = 246 | 0.47 [0.37, 0.57] | 0.098 [0.082, 0.114] | 0.093 [0.079, 0.109] | 0.11 [-0.08, 0.31] | 0.0087 [0.0075, 0.0097] |
| moderate FB *n* =421 | 0.54 [0.47, 0.61] | 0.074 [0.066, 0.083] | 0.071 [0.063, 0.079] | 0.12 [-0.03, 0.26] | 0.0063 [0.0056, 0.007] |
| UB *n* = 370 | 0.47 [0.4, 0.55] | 0.081 [0.07, 0.091] | 0.077 [0.068, 0.088] | 0.06 [-0.1, 0.21] | 0.0056 [0.0049, 0.0064] |
| moderate MB *n* =346 | 0.49 [0.4, 0.57] | 0.054 [0.046, 0.062] | 0.063 [0.055, 0.073] | 0.12 [-0.04, 0.28] | 0.0041 [0.0035, 0.0048] |
| high MB *n* = 167 | 0.37 [0.26, 0.49] | 0.072 [0.056, 0.088] | 0.048 [0.037, 0.062] | 0.24 [-0.05, 0.52] | 0.0026 [0.0019, 0.0034] |

continued

**Table S11** continued.

| | $F_{AI}$[1] | $\sigma_{AI,F}$[2] | $\sigma_{AI,M}$[3] | $\rho_{MF}$[4] | $\upsilon$[5] |
|---|---|---|---|---|---|
| **whole body (main cross)** | | | | | |
| All genes $n$ = 2170 | 0.43 [0.4, 0.44] | 0.063 [0.062, 0.066] | 0.052 [0.051, 0.054] | 0.64 [0.6, 0.66] | 0.0016 [0.0015, 0.0016] |
| high FB $n$ = 313 | 0.60 [0.51, 0.68] | 0.038 [0.033, 0.043] | 0.038 [0.033, 0.043] | 0.24 [0.1, 0.39] | 0.0013 [0.001, 0.0016] |
| moderate FB $n$ = 888 | 0.51 [0.45, 0.55] | 0.053 [0.049, 0.056] | 0.036 [0.033, 0.039] | 0.37 [0.27, 0.47] | 0.0014 [0.0012, 0.0016] |
| UB $n$ = 509 | 0.34 [0.27, 0.41] | 0.077 [0.066, 0.088] | 0.072 [0.063, 0.082] | 0.82 [0.75, 0.87] | 0.002 [0.0017, 0.0023] |
| moderate MB $n$ = 389 | 0.39 [0.32, 0.45] | 0.082 [0.073, 0.093] | 0.069 [0.060, 0.079] | 0.80 [0.74, 0.86] | 0.0016 [0.0013, 0.0019] |
| high MB $n$ = 77 | 0.47 [0.29, 0.65] | 0.050 [0.034, 0.067] | 0.044 [0.030, 0.059] | 0.92 [0.82, 1] | 0.0011 [0.0005, 0.0018] |
| **whole body (reciprocal cross)** | | | | | |
| All genes $n$ = 2386 | 0.35 [0.32, 0.37] | 0.069 [0.066, 0.073] | 0.060 [0.056, 0.064] | 0.67 [0.64, 0.69] | 0.0026 [0.0025, 0.0027] |
| high FB $n$ = 272 | 0.56 [0.45, 0.65] | 0.035 [0.029, 0.040] | 0.036 [0.03, 0.041] | 0.16 [-0.01, 0.35] | 0.0023 [0.0019, 0.0027] |
| moderate FB $n$ = 858 | 0.37 [0.31, 0.42] | 0.063 [0.056, 0.070] | 0.047 [0.043, 0.053] | 0.43 [0.34, 0.53] | 0.0028 [0.0025, 0.003] |
| UB $n$ =572 | 0.41 [0.36, 0.46] | 0.051 [0.045, 0.057] | 0.046 [0.041, 0.051] | 0.61 [0.5, 0.7] | 0.002 [0.0018, 0.0023] |
| moderate MB $n$ =530 | 0.34 [0.29, 0.39] | 0.097 [0.087, 0.109] | 0.090 [0.081, 0.102] | 0.84 [0.79, 0.88] | 0.0029 [0.0026, 0.0032] |
| high MB $n$ =154 | 0.43 [0.32, 0.55] | 0.057 [0.044, 0.071] | 0.047 [0.037, 0.059] | 0.94 [0.88, 1] | 0.0024 [0.0019, 0.0029] |

[1] $F_{AI}$ = frequency of genes with AI

[2] $\sigma_{AI,F}$ = average magnitude of AI in females; $\sigma_{AI,F}$ is the standard deviation of female effects, which is modelled as approximately normally distributed with mean 0

[3] $\sigma_{AI,M}$ = average magnitude of AI in males; $\sigma_{AI,M}$ is the standard deviation of male effects, which is modelled as approximately normally distributed with mean 0

[4] $\rho_{MF}$ = intersexual correlation in AI effects

[5] $\upsilon$ = overdispersion parameter

**Table S12.** Pairwise differences [with 95% high posterior density interval] in $F_{AI}$ between different categories of sex bias. Difference is (row category) – (column category) as estimated from model framework 2 (Table S11).

| | moderate FB | UB | moderate MB | high MB |
|---|---|---|---|---|
| heads | | | | |
| moderate FB | | **0.19**<br>**[0.13, 0.25]** | **0.33**<br>**[0.24, 0.42]** | NA |
| UB | | | **0.14**<br>**[0.06, 0.21]** | NA |
| gonads | | | | |
| hFB | -0.07<br>[-0.19, 0.05] | -0.01<br>[-0.13, 0.12] | -0.02<br>[-0.14, 0.11] | 0.1<br>[-0.06, 0.24] |
| mFB | | 0.06<br>[-0.04, 0.17] | 0.05<br>[-0.06, 0.16] | **0.16**<br>**[0.03, 0.3]** |
| UB | | | -0.01<br>[-0.13, 0.11] | 0.1<br>[-0.03, 0.24] |
| mMB | | | | 0.11<br>[-0.03, 0.25] |
| whole body (main cross) | | | | |
| | 0.09<br>[-0.01, 0.2] | **0.26**<br>**[0.13, 0.37]** | **0.21**<br>**[0.09, 0.32]** | 0.12<br>[-0.08, 0.32] |
| | | **0.16**<br>**[0.08, 0.25]** | **0.12**<br>**[0.04, 0.2]** | 0.03<br>[-0.16, 0.21] |
| | | | -0.05<br>[-0.14, 0.05] | -0.13<br>[-0.34, 0.05] |
| | | | | -0.09<br>[-0.28, 0.11] |
| whole body (reciprocal cross) | | | | |
| | **0.19**<br>**[0.07, 0.3]** | **0.15**<br>**[0.03, 0.26]** | **0.22**<br>**[0.1, 0.33]** | 0.13<br>[-0.04, 0.28] |
| | | -0.04<br>[-0.12, 0.03] | 0.03<br>[-0.05, 0.1] | -0.06<br>[-0.2, 0.07] |
| | | | 0.07<br>[-0.01, 0.14] | -0.02<br>[-0.15, 0.12] |
| | | | | -0.09<br>[-0.22, 0.04] |

**Table S13.** Parameter estimates [with 95% high posterior density interval] for the joint distribution of AI effects across heads and gonads using model framework 2 ("inter-tissue correlation of AI effects" model).

| | **Frequency of genes with AI,** $F_{AI}$ | **Average magnitude AI in heads**[1]**,** $\sigma_{AI,H}$ | **Average magnitude AI in gonads**[1]**,** $\sigma_{AI,G}$ | **Inter-tissue correlation of AI effects,** $\rho_{GHI}$ | **Overdispersion parameter,** $\upsilon$ |
|---|---|---|---|---|---|
| Female | 0.42 | 0.040 | 0.087 | 0.27 | 0.0037 |
| | [0.39, 0.45] | [0.037, 0.043] | [0.082, 0.092] | [0.19, 0.37] | [0.0035, 0.0039] |
| Male[2] | 0.43 | 0.049 | 0.090 | 0.21 | 0.0011 |
| | [0.41, 0.46] | [0.043, 0.052] | [0.083, 0.096] | [0.16, 0.27] | [0.0005, 0.0015] |

[1] the distribution of AI effects in each sex is modelled as approximately normally distributed with mean 0 and standard deviation $\sigma_{AI,t}$ in tissue $t$

[2] A modified procedure was used to approximate the posterior distributions for males because the MCMC chain was ill-formed following the 'standard' procedure used in other analyses because almost all proposed values during MCMC were rejected. In principle, this could be resolved by running much longer chains but this would be computationally prohibitive due to the time needed to calculate each likelihood. We approximated posterior using the following procedure. We created a list of all parameter values proposed during the standard procedure and their likelihoods. We then performed an MCMC-like procedure on this list (i.e., proposing values from this list and accepting or rejecting these following standard MCMC criteria). This is computationally many times faster because the likelihoods are known and, thus, allows much longer chains to be run. We tested this procedure with other data sets where this procedure was not necessary (e.g., the female data) and obtained similar values to the standard procedure.

Figure S1.

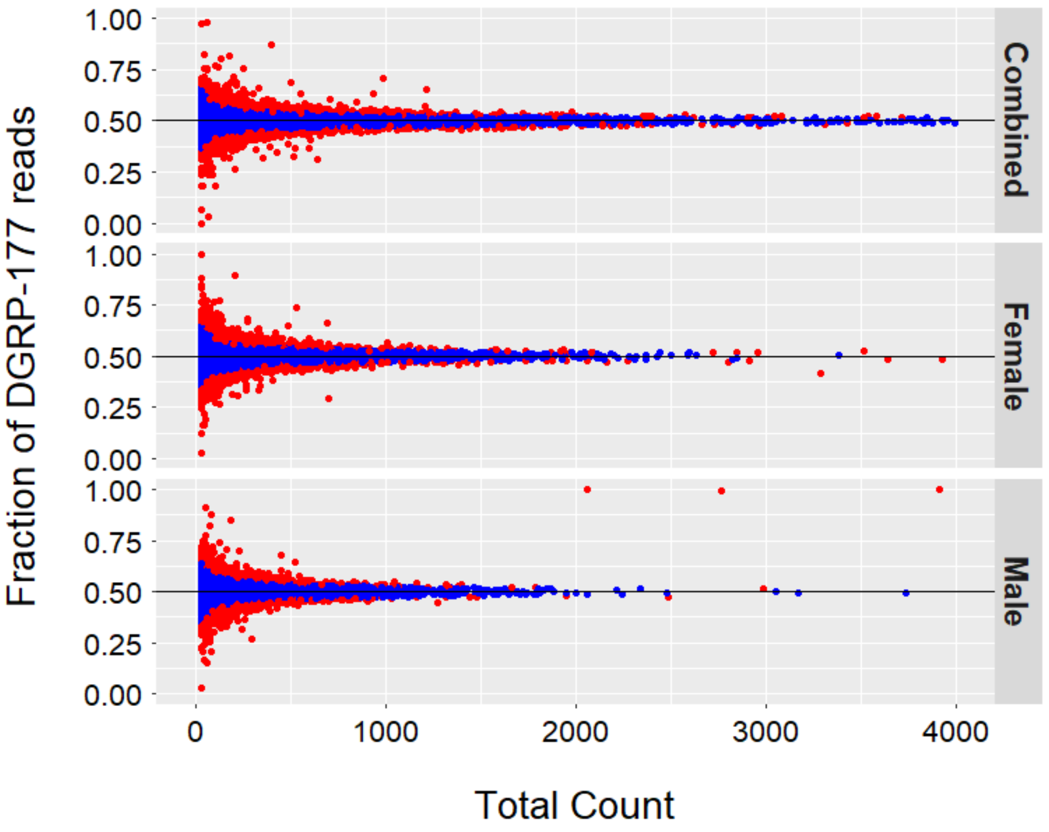


**Figure S1**. Assignment of F1 genomic DNA reads. The fraction of DGRP-177 reads in genomic data tends to be around 0.5. Genes that deviate substantially from this expected proportion (shown in red) are marked for filtering due to potential mapping bias. In the absence of mapping bias, the ratio of allelic reads in genomic data should ideally be 1:1, and accordingly, read counts were assumed to follow a binomial distribution with probability of being DGRP-177 of 0.5. If the binomial probability of the observed DGRP-177 read counts for a given fell within either of the 5% tails, the gene was marked as having potential evidence of mapping bias (red points). Thus, by this method, we expect 10% of genes with no true bias to fall into these tails (i.e., false positives for mapping bias). Genes were tested separately using male and female data, and also using combined data from the sexes. 1681 genes, out of 12045 genes (14% of genes) in the combined data, had evidence of mapping bias. 1560 genes in females (13% of genes), and 1448 genes in males (12% of genes) had evidence of the same. All the 3102 genes (of 12045 genes; 25%) were excluded from further analysis. We also excluded genes where coverage of DGRP-177 reads deviates from the expected proportion of 0.5 by more than 0.1 in either male or female data. 1184 genes (of 12045 genes), many of which also failed previous criteria, had significant deviation from this expectation. As a result, an additional 197 genes were excluded from analysis.

Figure S2

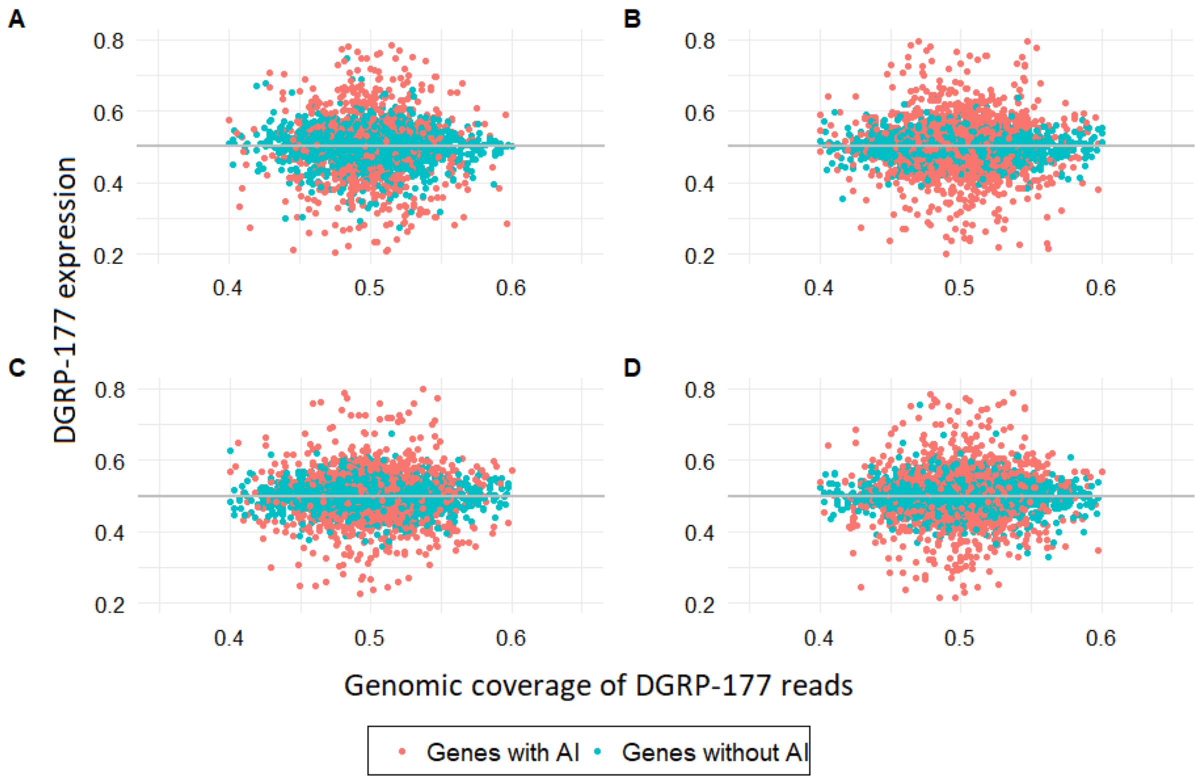


**Figure S2**. Allele frequencies estimated from genomic vs. transcriptomic reads from male F1 samples. These analyses are using the set of genes post-filtering, i.e., the set of genes for the results shown in Table 1. Fraction of DGRP-177 reads in male F1 genomic data versus fraction of expression of DGRP-177 in RNA-seq for A) gonads, B) heads, C) whole bodies (main cross), and D) whole bodies (reciprocal cross). As shown here and in Table S4, the correlation of DGRP-177 expression and DGRP-177 genomic coverage is nearly zero for all samples. This indicates that the filtering criteria is largely effective in mitigating instances of mapping bias in the data.

Figure S3

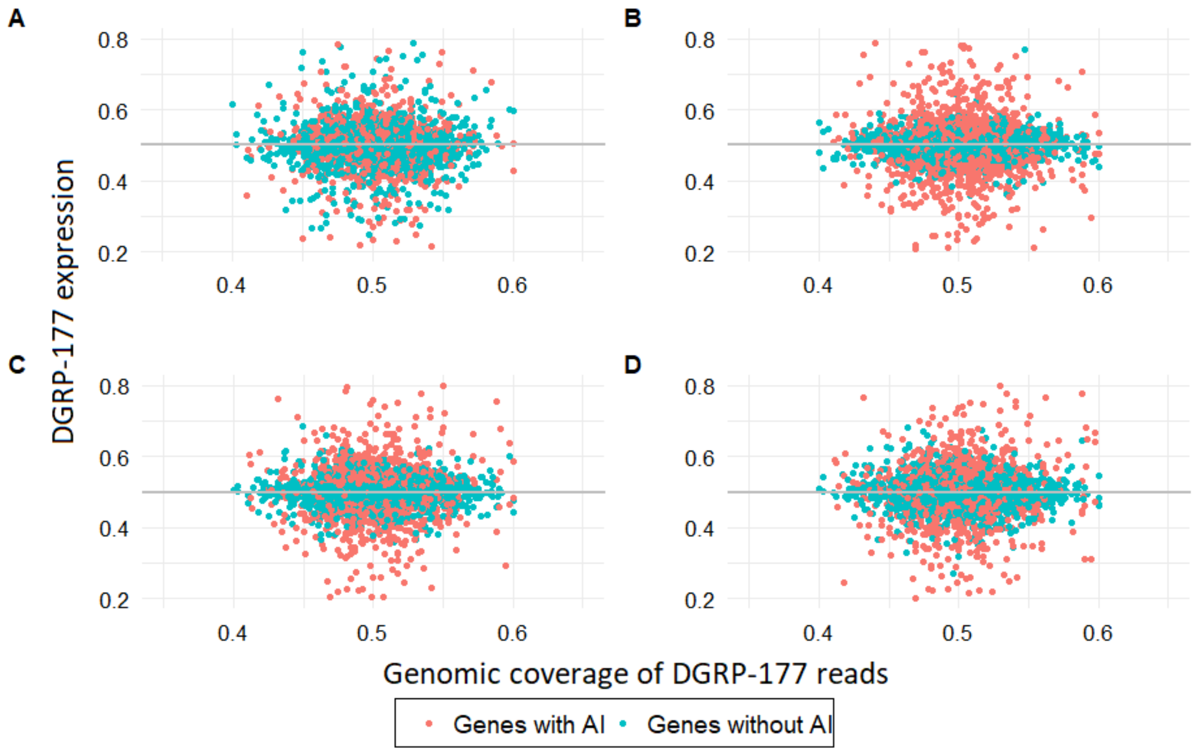


**Figure S3**. Allele frequencies estimated from genomic vs. transcriptomic reads from female F1 samples. These analyses are using the set of genes post-filtering, i.e., the set of genes for the results shown in Table 1. These are the genes for the analyses shown in Table 1. Fraction of DGRP-177 reads in male F1 genomic data versus fraction of expression of DGRP-177 in RNA-seq for A) gonads, B) heads, C) whole bodies (main cross), and D) whole bodies (reciprocal cross). As shown here and in Table S5, the correlation of DGRP-177 expression and DGRP-177 genomic coverage is nearly zero for all samples. This indicates that the filtering criteria is largely effective in mitigating instances of mapping bias in the data.

Figure S4

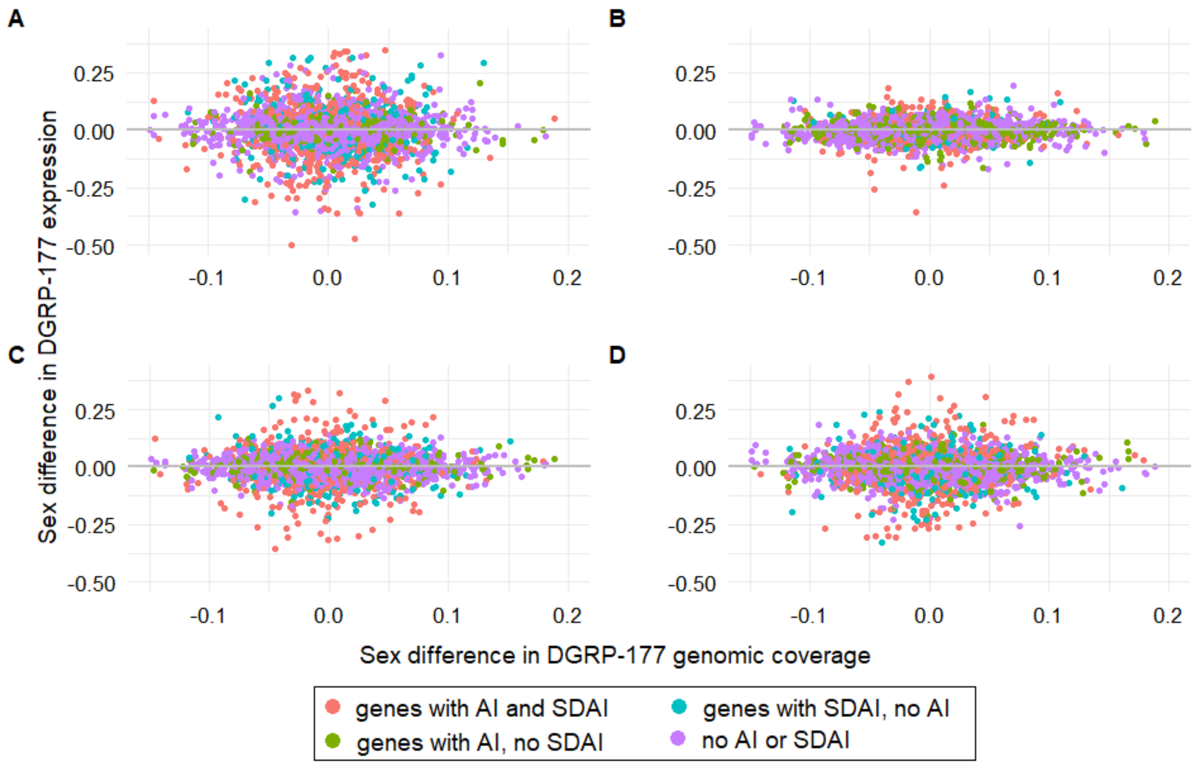


**Figure S4**. Sex differences in allele frequency estimated from genomic vs. transcriptomic reads from F1 samples. These analyses are using the set of genes post-filtering, i.e., the set of genes for the results shown in Table 1. The sex difference in the fraction of DGRP-177 reads in genomic DNA vs in the fraction of DGRP-177 reads in expression in F1 for A) gonads, B) heads, C) whole bodies (main cross), D) whole bodies (reciprocal cross).

Figure S5

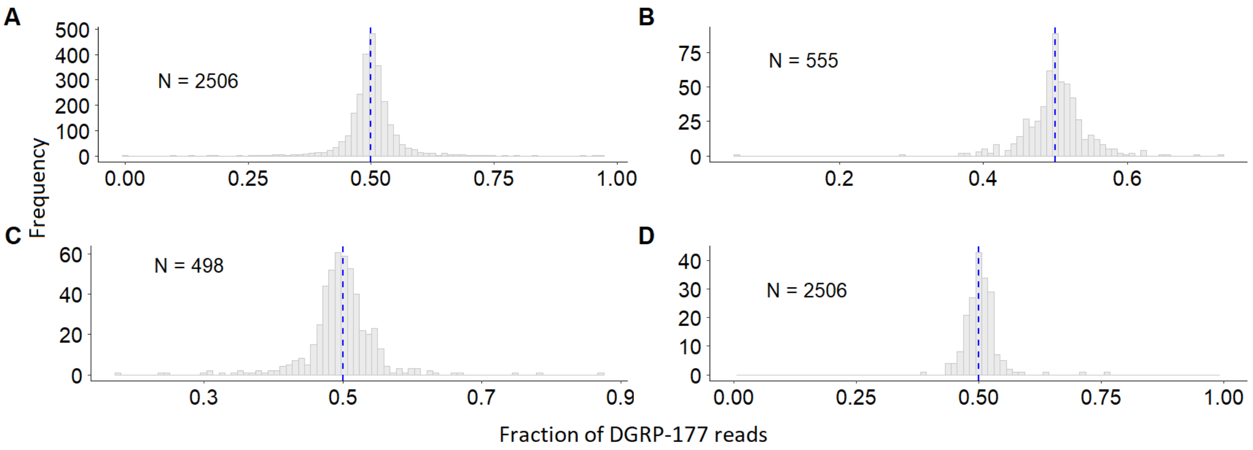


**Figure S5.** Distribution of the estimated frequency of DGRP-177 RNA-seq reads from whole body F1 samples, stratified with respect to parental genomic DNA read assignment accuracy. Frequency of DGRP-177 reads in A) genes with >99% correctly-mapped reads in both parents (n = 2133), B) genes with >99% correctly-mapped reads in DGRP-177, but not SP159N (n = 459), C) genes with >99% correctly-mapped reads in SP159N, but not DGRP-177 (n = 419), D) genes with <99% correctly-mapped reads in both parents (n = 166). The blue-dashed vertical line corresponds to the point where fraction of DGRP-177 reads is $f = 0.5$.

Figure S6

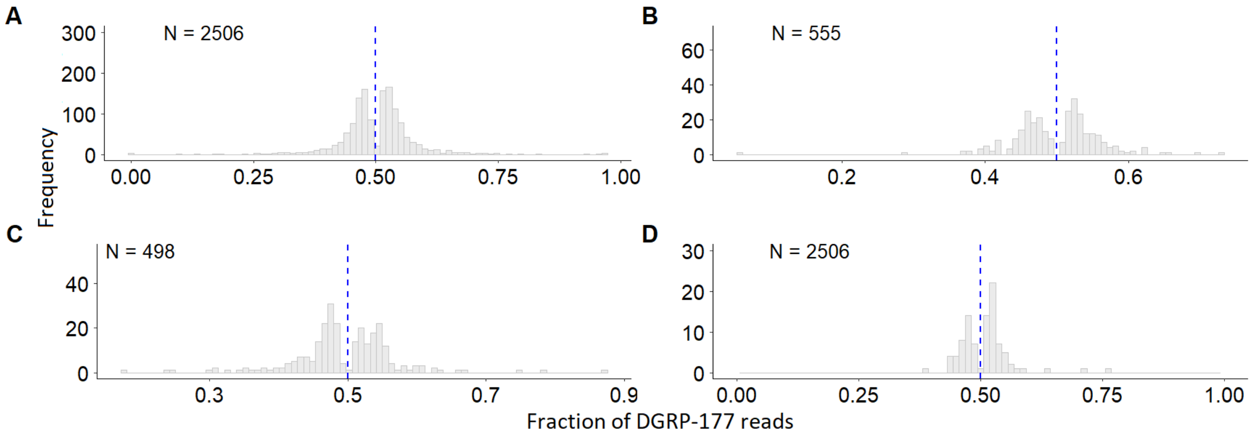


**Figure S6.** Distribution of the estimated frequency of DGRP-177 RNA-seq reads from whole body F1 samples using only genes with significant AI, stratified with respect to parental genomic DNA read assignment accuracy. Frequency of DGRP-177 reads in A) genes with >99% correctly-mapped reads in both parents (n = 1225), B) genes with >99% correctly-mapped reads in DGRP-177, but not SP159N (n = 240), C) genes with >99% correctly-mapped reads in SP159N, but not DGRP-177 (n = 244), D) genes with <99% correctly-mapped reads in both parents (n = 82). The blue-dashed vertical line corresponds to the point where fraction of DGRP-177 reads is $f = 0.5$.
